# Removal of extracellular human amyloid beta aggregates by extracellular proteases in *C. elegans*

**DOI:** 10.1101/2022.09.14.507993

**Authors:** Elisabeth Jongsma, José María Mateos, Collin Y. Ewald

**Author notes:** Corresponding authors (CYE).

## Abstract

The amyloid-beta (Aβ) plaques found in Alzheimer’s disease (AD) patients’ brains contain collagens and are embedded extracellularly. Several collagens have been proposed to influence Aβ aggregate formation, yet their role in clearance is unknown. To investigate the potential role of collagens in forming and clearance extracellular aggregates *in vivo*, we created a transgenic *Caenorhabditis elegans* strain that expresses and secretes human Aβ_1-42_. This secreted Aβ forms aggregates in two distinct places within the extracellular matrix. In a screen for extracellular human Aβ aggregation regulators, we identified different collagens to ameliorate or potentiate Aβ aggregation. We show that a disintegrin and metalloprotease ADM-2, an orthologue of ADAM9, reduces the load of extracellular Aβ aggregates. ADM-2 is required and sufficient to remove the extracellular Aβ aggregates. Thus, we provide *in-vivo* evidence of collagens essential for aggregate formation and metalloprotease participating in extracellular Aβ aggregate removal.

**Highlights:** Extracellular aggregates of amyloid beta are a hallmark of Alzheimer’s disease. Here we developed a novel *C. elegans* transgenic line that secretes human amyloid beta, which forms aggregates in the extracellular matrix (ECM). We show that ECM dynamics can disturb aggregation and that ADM-2, an ortholog of Human ADAM9, is involved in removing these extracellular aggregates.

## Introduction

Alzheimer’s disease currently affects >1 in 9 people above 65 years of age (11.2%) in the USA and is the seventh cause of death worldwide (Alzheimer’s Association, 2021.; Mortality and Global Health Estimates, 2019). A hallmark of Alzheimer’s disease is the extracellular aggregation of amyloid-beta. Over the past decades, much knowledge has been gained on the production and removal of amyloidbeta (Aβ). Several mechanisms are involved in clearing Aβ from the brain, including enzymatic and non-enzymatic pathways. The non-enzymatic pathways include the continuous flow of the interstitial fluid into the cerebrospinal fluid followed by interstitial fluid drainage, phagocytosis by microglia or astrocytes, and receptor-mediated transport across the blood-brain barrier (Elbert et al., 2022; Sagare et al., 2007; Tajbakhsh et al., 2021; Zhao et al., 2015). The enzymatic pathway involves several proteases, including matrix metalloproteinases, neprilysin, insulin-degrading enzymes, and glutamate carboxypeptidase. The extracellular protein heparan sulfate proteoglycans (HSPGs) can block the clearance of Aβ. HSPGs are often found in Aβ depositions where they might block enzymatic degradation (Gupta-Bansal et al., 1995; Su et al., 1992; van Horssen et al., 2003). Moreover, while most HSPGs promote the uptake of Aβ through lipid rafts, uptake of Aβ through clathrin-mediated endocytosis is blocked when the HSPG (SDC3) binds to Aβ (Letoha et al., 2019).

One of the least understood observations is the consistent co-aggregation of specific collagens with Aβ plaques. Interestingly, the compaction of Aβ into plaques can be influenced by the expression of collagenous amyloid plaque components (CLACs) (Hashimoto et al., 2020). CLAC is a collagen type XXV a1 chain (COL25A1) cleavage product. COL25A1 overexpression can have detrimental effects in mice (Tong et al., 2010). However, human genetic studies suggest a more complex interplay where certain single nucleotide polymorphisms in COL25A1 are associated with AD and others are, in contrast, associated with health effects in the elderly (Erikson et al., 2016; Forsell et al., 2010). Curiously, several other collagens have been found to have a protective role. Co-localizing with vascular amyloid at the basal lamina is collagen XVIII, a heparan sulfate proteoglycan that reduces disease symptoms (Van Horssen et al., 2002). Collagen VI was found at the ECM and the basal lamina and can block the interaction between neurons and oligomers and help protect against neurotoxicity (Cheng et al., 2009; Ma et al., 2020). Furthermore, the basement lamina collagen IV was 55% upregulated in cerebral vessels when comparing AD to healthy subjects (Cheng et al., 2009; Farkas et al., 2000; Kalaria & Pax, 1995; Nguyen et al., 2021). This upregulation is specific to the brain region and Braak stage (Lepelletier et al., 2017). Collagen IV was shown to bind the amyloid precursor protein (APP), prevent Aβ fibril formation, and even disrupt preformed Aβ fibrils (Kiuchi, Isobe, & Fukushima, 2002; Kiuchi, Isobe, Fukushima, et al., 2002; Narindrasorasak et al., 1995). Based on these collected studies, we hypothesized that the ECM may be more than a passive bystander and that its components hold the potential to influence disease progression.

To address this hypothesis, we set out to explore the mechanisms by which ECM components influence amyloid beta aggregate formation and clearance *in vivo*. However, a model monitoring this *in-vivo* and non-invasively was missing. Therefore, we generated a novel transgenic *C. elegans* strain, with inducible expression and secretion of human Aβ_1-42_ tagged with super-folder GFP (sfGFP::Aβ). Furthermore, *C. elegans* is a suitable model to address this question. Several Alzheimer-related pathways are highly conserved between humans and *C. elegans* (Apostolakou et al., 2021; Ewald & Li, 2010). Moreover, the ECM components associated with AD have orthologs in *C. elegans*. The *C. elegans* EMB-9 and LET-2 are collagen type IV, CLE-1 is collagen type XVIII, and COL-99 is collagen type XXV (Teuscher, Jongsma, et al., 2019). While EMB-9 and LET-2 localize to the basal lamina, CLE-1 and COL-99 localize to neurons.

Here, we show that upon induction, secreted sfGFP::Aβ is initially cleared by the excretory system, the gut, and the coelomocytes. However, Aβ is retained past 24h and forms non-mobile structures in the ECM. We identified collagens that can completely suppress Aβ aggregate formation. Moreover, we find modulators of the ECM, metalloproteases, to assist in the removal of extracellular Aβ aggregates. We demonstrate that one of these metalloproteases, ADM-2 is essential to remove Aβ aggregates. Taken together, this suggests that ECM composition is critical to allow Aβ aggregate formation, while dynamic regulation of the ECM through metalloproteases is key in Aβ aggregate clearance.

## Results

### Generating an *in-vivo* model for extracellular Aβ aggregates

An obstacle to studying the interaction of the ECM with amyloid-beta aggregation and clearance is the lack of an *in-vivo* model. Previous human Aβ expressing *C. elegans* strains failed to secrete Aβ and model intracellular Aβ toxicity (Ewald & Li, 2012; Link, 1995). Therefore, we designed a genetic construct that secretes Aβ tagged with GFP (Figure 1A, 1B). Expression of this construct was induced by heat shock under the control of the *hsp-16.2* promoter that drives expression in many tissues but predominantly in neurons and hypodermis (Bacaj & Shaham, 2007). Furthermore, the construct has a longer 3’ UTR targeting its mRNA for non-sense mediated degradation to prevent the leakage of the *hsp-16.2* promoter (Ewald et al., 2016). This allowed us to separate events scaled in time, for example, deposition versus removal of Aβ. We used super-folder GFP (sfGFP) because it is more stable in the extracellular space than classical GFP (Pédelacq et al., 2006). A spacer sequence was placed between the sfGFP and the Aβ to allow the comparably smaller-sized Aβ to move and interact freely to form aggregates (Figure 1A, 1B). The full length of the Aβ_1-42_ peptide is essential for its aggregation (Mccoll et al., 2012). In our construct, Aβ is preceded by the sfGFP and spacer sequence, which prevents the truncation of the first few amino acids observed in many previous *C. elegans* Aβ models (McColl et al., 2009). As controls, we generated two constructs; one containing a non-aggregating version of Aβ_1-42_ (secreted sfGFP::Aβ(F20S, L35P)) (Wurth et al., 2002), and the other control is the secreted sfGFP without amyloid-beta fragment (Figure 1A).

**Figure 1.**
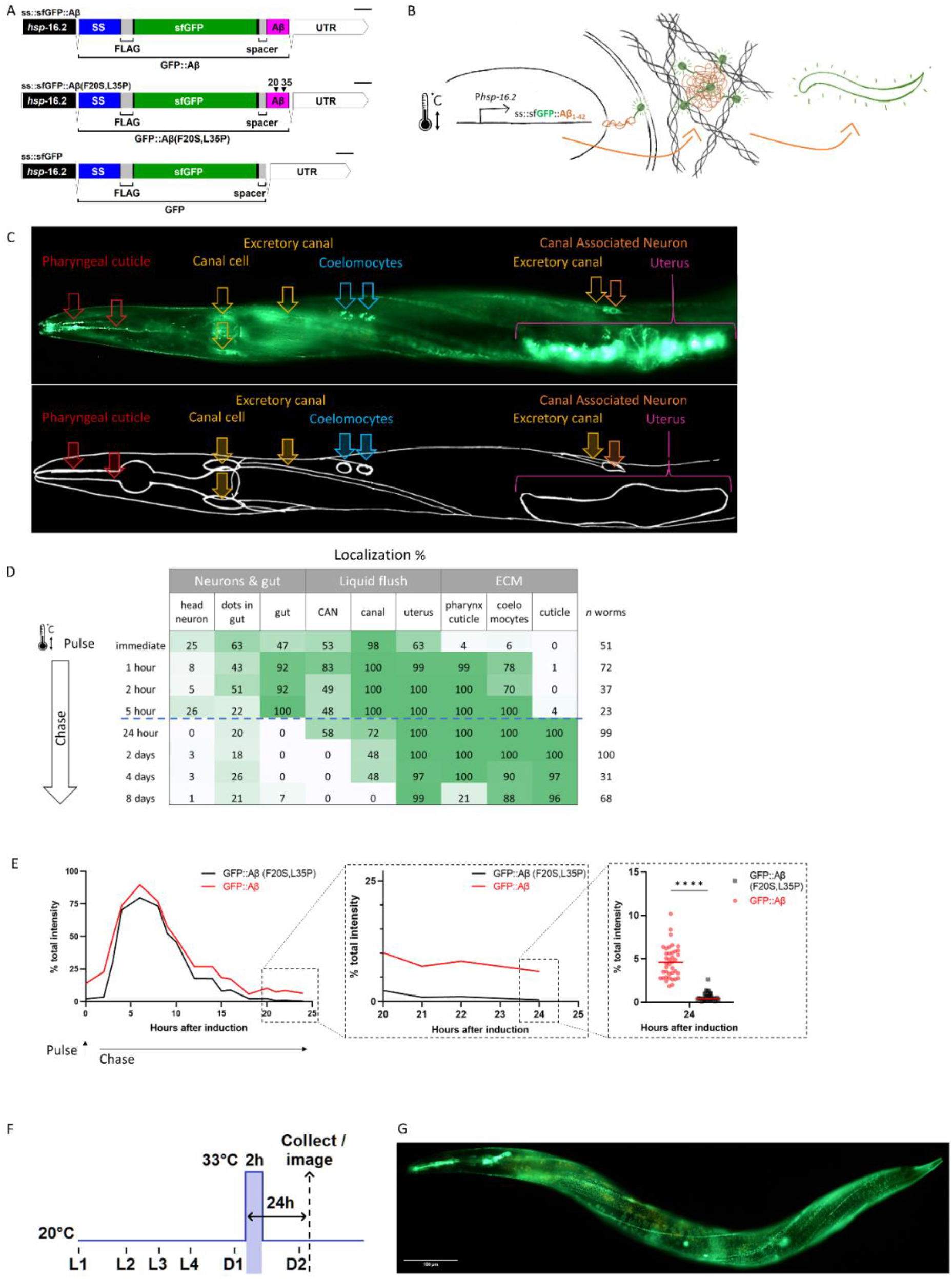
Expression of secreted human amyloid beta tagged with super-folder GFP. **A)** Genetic constructs used to generate transgenic *C. elegans* strains. SS indicates secretion sequence, UTR = untranslated region **B)** Hypothetical model of induction, expression, and secretion of sfGFP::Aβ in *C. elegans*. After a single heat shock induction, sfGFP::Aβ is secreted and localizes to different tissues over time. **C)** Localization of sfGFP::Aβ to different tissues in *C. elegans*. The localization to the excretory canal and the coelomocytes confirm that the sfGFP::Aβ is secreted. **D)** Percentage of tissue type with sfGFP::Aβ over time. After production and secretion, most of the produced sfGFP::Aβ was flushed out, but some was retained at the cuticle up to 8 days after the induction event. **E)** Clearance of sfGFP::Aβ is significantly slowed >18h after induction compared to non-aggregating control sfGFP::Aβ(F20S, L35P). Data represented is the average from three independent repeats combined; repeats are shown in supporting figure 1E. Figure 1E, sub III image is from one of the repeats, unpaired, two-tailed t-test. ****: p < 0.0001. **F)** Representation of methods regarding the time of heat shock induction of expression and imaging or sample collection 24h after induction. **G)** Representative image of the transgenic line LSD2104 and localization of secreted sfGFP::Aβ.

### Dynamic turnover of secreted amyloid beta

After a single heat shock induction of sfGFP::Aβ (*i.e*., single pulse-chase), sfGFP::Aβ was expressed in many tissues and secreted into the extracellular space surrounding different tissues (Figure 1C). This Aβ localization changed over time. Initially, Aβ localized to neurons in the nerve ring, the gut, the canal cells, the excretory canal, the canal-associated neurons, and the uterus (Figure 1C, 1D). During the first 24h, most of the sfGFP::Aβ appeared diffuse. After 24h, we observed bright puncta localized to the pharyngeal cuticle, the coelomocytes, and the cuticle (Figure 1D). Remarkably, the sfGFP::Aβ remained for an exceptionally long time at the cuticle, coelomocytes, and the uterus, up to eight days after induction (Figure 1D).

To determine whether this is due to the potential aggregation of Aβ, we followed sfGFP::Aβ intensity in a pulse-chase time course, including our two controls. After heat shock, all three strains showed similar time course trajectories for the induction (Supporting figure 1E.1). However, at the 24h time point where the sfGFP::Aβ was observed at the cuticle, the non-aggregating sfGFP::Aβ(F20S, L35P) and the sfGFP-only strains almost completely lost the GFP signal (Figure 1E, Supporting Figure 1E.1 and 1E.2). No localization to the cuticle was observed at this time point. Taken together, this suggests these sfGFP-linked Aβ were secreted and efficiently cleared from the extracellular spaces. At the same time, the retention at the cuticle after 24h was specific to Aβ_1-42_ due to its ability to form aggregates. In search of mechanisms selectively affecting the aggregation-prone amyloid-beta, all further interventions were assessed at the 24h (Figure 1F). We have generated a transgenic strain expressing and secreting aggregation-prone sfGFP::Aβ that, after initial clearance, is retained at the cuticle (Figure 1G).

### sfGFP::Aβ forms ‘flower’ and ‘moss’ patterns in the extracellular space

To characterize the potential sfGFP::Aβ aggregates in the extracellular space, we compared the non-aggregating sfGFP::Aβ(F20S, L35P) with the aggregation-prone sfGFP::Aβ. At 16 hours post-induction, the non-aggregating sfGFP::Aβ(F20S, L35P) formed a striped pattern, equally distributed along the body, reminiscent of the striped pattern of furrows on the cuticle. This signal was relatively weak and uniform, similarly to the GFP-only strain (Figure 2A (intensity enhanced), Supporting figure2.1). By contrast, sfGFP::Aβ formed two types of bright structures, which we named ‘flower’ and ‘moss’ (Figure 2B). Interestingly, the ‘flower’ structures were exclusively found over the nematode’s dorsal and ventral sides, where muscle tissue underlies the hypodermis. The ‘moss’ structures were consistently found on the left and right sides of the nematode, apical to the main hypodermal syncytium (hyp7). The localization of the aggregate type is invariant and correlates with the underlying tissue type. The flower and moss structures are not found for the non-aggregating sfGFP::Aβ(F20S, L35P) (Figure 2A) nor for GFP-only (Supporting figure 2.1) but were unique to the wild-type sfGFP::Aβ strain.

**Figure 2.**
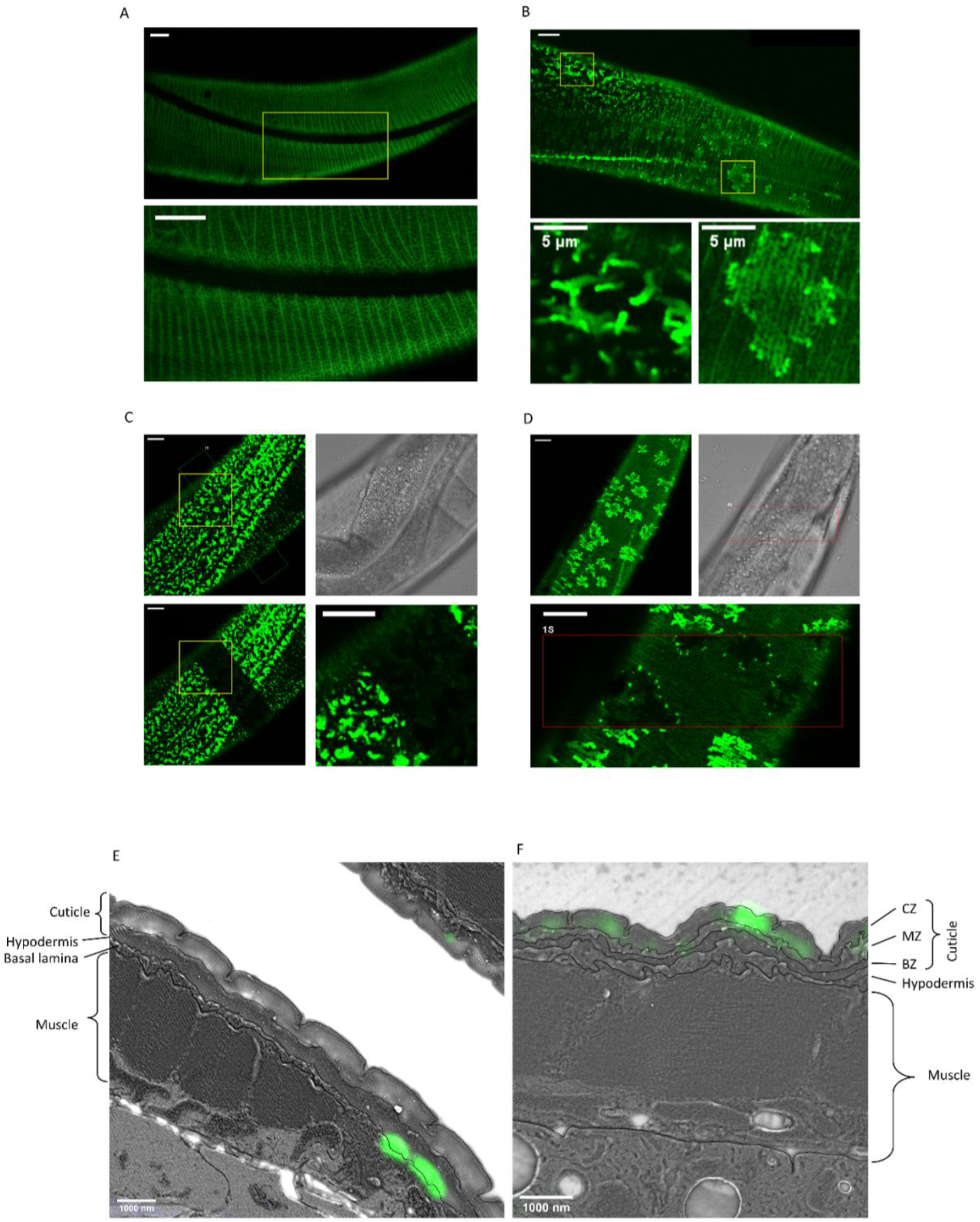
Secreted amyloid beta form aggregates at *C. elegans* ECM. **A)** The control strain sfGFP::Aβ(F20S, L35P) shows localization near the cuticle. However, the signal is relatively weak and uniform. Note. Image intensity enhanced, taken 16h after induction. **B)** Two types of bright patterns, dubbed ‘moss’ and ‘flower’, can be observed for sfGFP::Aβ near the cuticle, 24h past induction of expression. **C, D)** Fluorescence recovery after photobleaching shows both the moss (**C**) and flower (**D**) structures are immobile. Time of imaging up to 4h after bleaching. **E, F)** Correlative light electron microscopy revealed localization to ECM structures. **E)** Localization of the “moss” structures to basal lamina when there is no muscle underneath. **F)** Localization of the “flower” structures to the cuticle when there is muscle underneath. CZ: cortical zone of the cuticle, MZ: medial zone of the cuticle, BZ: basal zone of the cuticle. Scale bars are 10 μm unless otherwise indicated.

### The ‘Flower’ and ‘moss’ patterns are immobile aggregates

Next, to determine whether these flower and moss structures could be aggregates, we used fluorescence recovery after photobleaching (FRAP), a technique commonly used to determine if the tagged protein of interest is fixed in place. A small area within the image field was photobleached using a brief exposure to UV, quenching the GFP fluorescence. Free movement or transport of the fluorescently-tagged protein should lead to an exchange of molecules between the bleached- and unbleached area, resulting in a recovery of fluorescence in the bleached (dark) area over time. In the moss structure, no such recovery was observed (Figure 2C). Even when one-half of a single moss structure was photobleached, there was no recovery in the bleached area, indicating no movement or exchange of molecules even within the structure (Figure 2C). Upon photobleaching, the flower structures turn completely dark (Figure 2D). The surrounding, relatively low GFP intensity recovered rapidly (within seconds), but the flower structure remained dark. No recovery of signal inside the structure was observed; however, after four hours, some bright dots were observed on the edge of the flower structures, indicating potential growth or exchange of Aβ from outside the photobleached area. However, this was observed only at the border and not at the center of the flower structures (Figure 2D), suggesting confinement on the upper and lower part of this structure as it would be “sandwiched” between something. For both the flower and moss structures, the absence of fluorescence recovery for up to four hours confirmed that these structures are immobile, suggesting that these structures are aggregates.

### Distinct aggregate patterns on the cuticular extracellular matrix

To determine where these structures localize, we used correlative light and electron microscopy (Mateos et al., 2018). This technique localizes the fluorescence signal in relation to the morphology of the tissue on the same thin sections (110 nm), providing a high X, Y, and Z resolution. The sfGFP::Aβ was observed to localize to two different parts of the cuticular extracellular matrix (Figure 2E, 2F, Supporting figures 2.2–2.4). The localization apical to the hypodermis, underneath the collagen-dense cuticle, is representative of the ‘moss’ type aggregates since these are associated with the hypodermis (Figure 2E, Supporting figures 2.2,2.3). By contrast, the localization to the cortical layer of the cuticle is representative of the ‘flower’ type aggregates since these are associated with underlying muscle tissue (Figure 2F, Supporting figure 2.4). Taken together, we established a novel *in-vivo* model of secreted human Aβ that forms aggregates in the extracellular matrix.

### Screening identified clearance mechanisms of Aβ aggregates in the extracellular matrix

To identify key molecular players in the formation and clearance of Aβ aggregates localized in the extracellular matrix (ECM), we designed a targeted RNA interference (RNAi) screen. We rationalized that examining four main categories of genes could elucidate a role for ECM molecules in Aβ aggregation. These categories were: all known ECM genes and ECM remodeling genes (matrisome; n=719)(Teuscher, Jongsma, et al., 2019), genes involved in attaching the cell to the ECM and mechanosensation (n=255), *C. elegans* orthologs of genes associated with human Alzheimer’s Disease (n=776) (Vahdati Nia et al., 2017) as well as genes protecting against neurodegenerative and age-related pathologies (n=631) (Figure 3A, Supplementary file 1.

**Figure 3.**
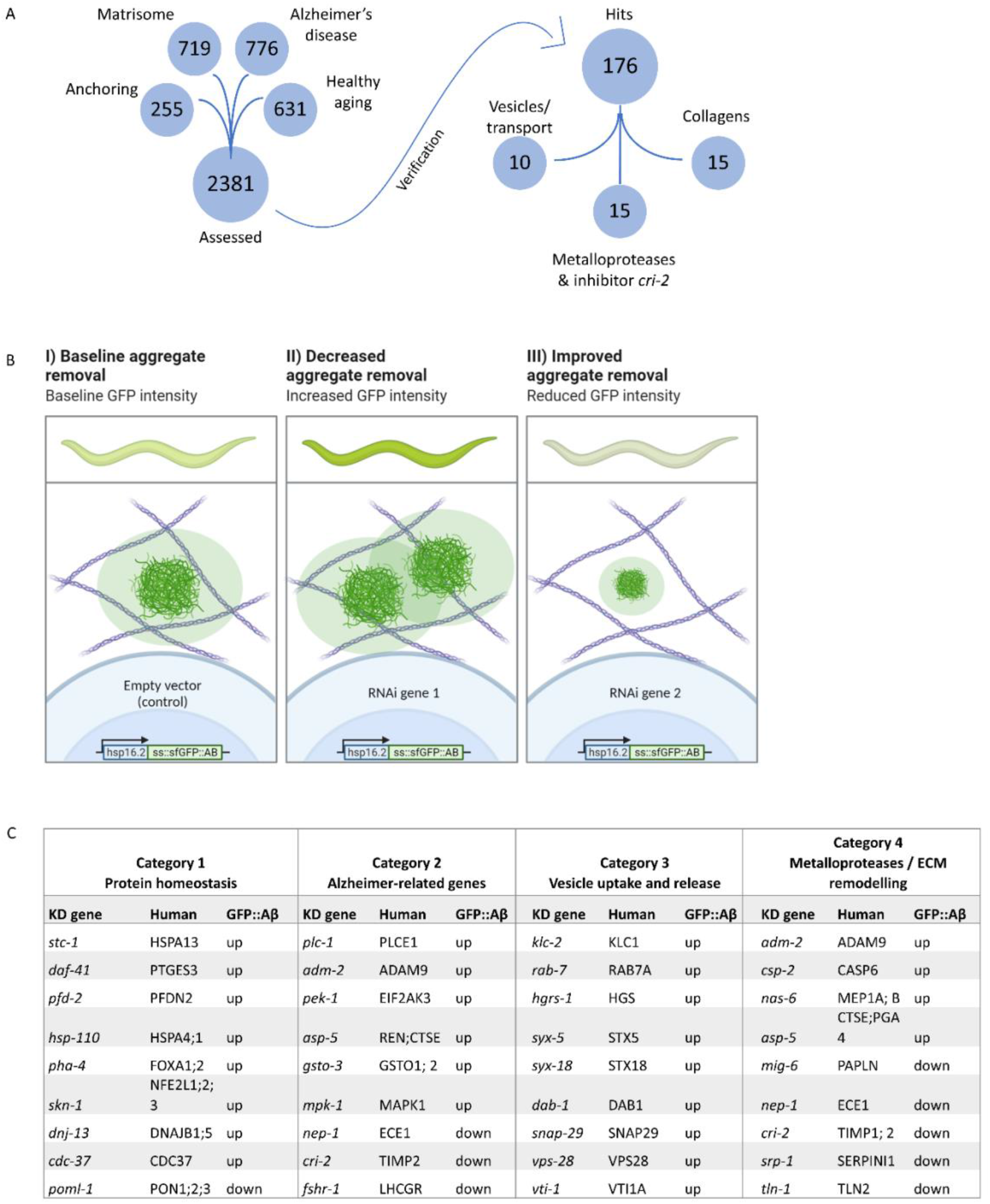
Strategy for RNA interference screening identified genes involved in the removal of extracellular sfGFP::Aβ. **A)** Four RNAi libraries were designed based on their hypothesized potential to affect extracellular sfGFP::Aβ aggregation. The Matrisome library contains all extracellular *C. elegans* genes, the Anchoring library contains transmembrane genes, the Alzheimer’s disease library is based on a meta-analysis of *C. elegans* orthologs of human GWAS studies, and the healthy aging library consists of genes that have a protective role against aging-related disease. Of the 2381 genes assessed, 176 genes were found to either increase- or decrease sfGFP::Aβ load. **B)** Expected fluorescence phenotypes of suppressor or enhancer genes. Grown on individual RNA clones from the L1 larval stage, populations of about 45 animals were assessed for an increase- or decrease in GFP signal. An increase of signal would indicate decreased aggregate removal, while a decrease of GFP signal would imply improved aggregate removal upon knockdown of target gene expression. **C)** Summary of categorization of hits shows relevant mechanisms to extracellular sfGFP::Aβ aggregate removal. Category 1 revealed screen hits in protein homeostasis, as expected when observing protein expression and turnover. Category 2 showed screen hits of orthologues of genes well known to be associated with Alzheimer’s disease in humans. Category 3 revealed screen hits of genes involved in vesicle uptake and release, processes essential to secretion and removal of extracellular proteins such as sfGFP::Aβ. Category 4 showed the involvement of metalloproteases and an inhibitor of metalloproteases. Selected candidate hits are shown; the full table and raw RNAi score are available in Supplementary file 2.

To assess the effects of individual gene knockdown on aggregation in the ECM, an increase or decrease in sfGFP::Aβ signal was scored 24h post-induction (Figure 3B), the time point at which non-aggregating Aβ and soluble GFP were flushed out. We identified 176 from 2368 screened genes to increase- or decrease sfGFP::Aβ intensity *in vivo* (Figure 3A; Supplementary file 1). To prioritize hits, we categorized these candidate genes into four groups (Figure 3C, Supplementary file 2). In category 1, we grouped hits expected to affect sfGFP::Aβ, such as genes involved in protein homeostasis, chaperones, and protein degradation (Figure 3C, Supplementary file 2). Reassuringly, we identified 71 orthologs of Alzheimer’s Disease implicated genes (category 2, Figure 3C, Supplementary file 2), suggesting the conservation of key molecular players relevant to human disease. We also identified interesting hits with less established relationships to Alzheimer’s disease, such as vesicle transport and vesicle fusion, as well as members of the SNARE complex (category 3, Figure 3C, Supplementary file 2). When interrogating these hits with follow-up experiments, the change in sfGFP::Aβ aggregation could be explained by alternative mechanisms, such as impaired secretion or endosome-recycling. Lastly, in category 4, we found that the knockdown of metalloproteinases (MMP) increased Aβ levels, whereas the knockdown of their inhibitors (TIMP) decreased the Aβ levels (Figure 3C, Supplementary file 2). Furthermore, knockdown of several individual collagens either increased or decreased sfGFP::Aβ fluorescence, which reinforced the idea of a potentially active role of ECM components in Aβ aggregation and Aβ aggregate removal (Figure 3C, Supplementary file 2).

### Collagens implicated in Aβ aggregate formation

To determine the role of collagens in Aβ aggregation and Aβ aggregate removal, we examined flower and moss Aβ aggregates upon collagen knockdown. We found that knockdown of cuticular collagen *col-2* or *col-79* resulted in more animals with flower Aβ aggregates (Figure 4A). By contrast, cuticular collagens *dpy-3* and *col-89* knockdown resulted in the complete absence of moss and flower Aβ aggregates, and *col-8*(RNAi) showed a marked reduction in Aβ aggregates (Figure 4A). As previously reported, DPY-3 is required for furrow formation of the cuticle, and animals lacking DPY-3 show disturbance of cuticular organization combined with a shortened and thicker body shape (Sandhu et al., 2021). DPY-3 is expressed during early developmental stages, while COL-8 and COL-89 are expressed during the last larval stage L4. The aggregates may require a particular ECM composition or specific collagens to form aggregates, or the knockdown of some collagens triggers overall ECM remodeling aiding the removal of aggregates. To separate these possibilities, we compared the knockdown of these collagens starting at different time points: RNAi beginning at the first larval stage (L1) to RNAi starting at the last larval stage (L4). We found that DPY-3 was required from early development for aggregates to form but not at later stages (Figure 4B, 4C). For COL-8 and COL-89, the effects of RNAi on intensity and aggregates were present when knocked down from the L4 stage (Figure 4B, 4C). Indeed, this suggests that the presence of these structural components of the ECM is either directly or indirectly required for Aβ aggregate formation.

**Figure 4.**
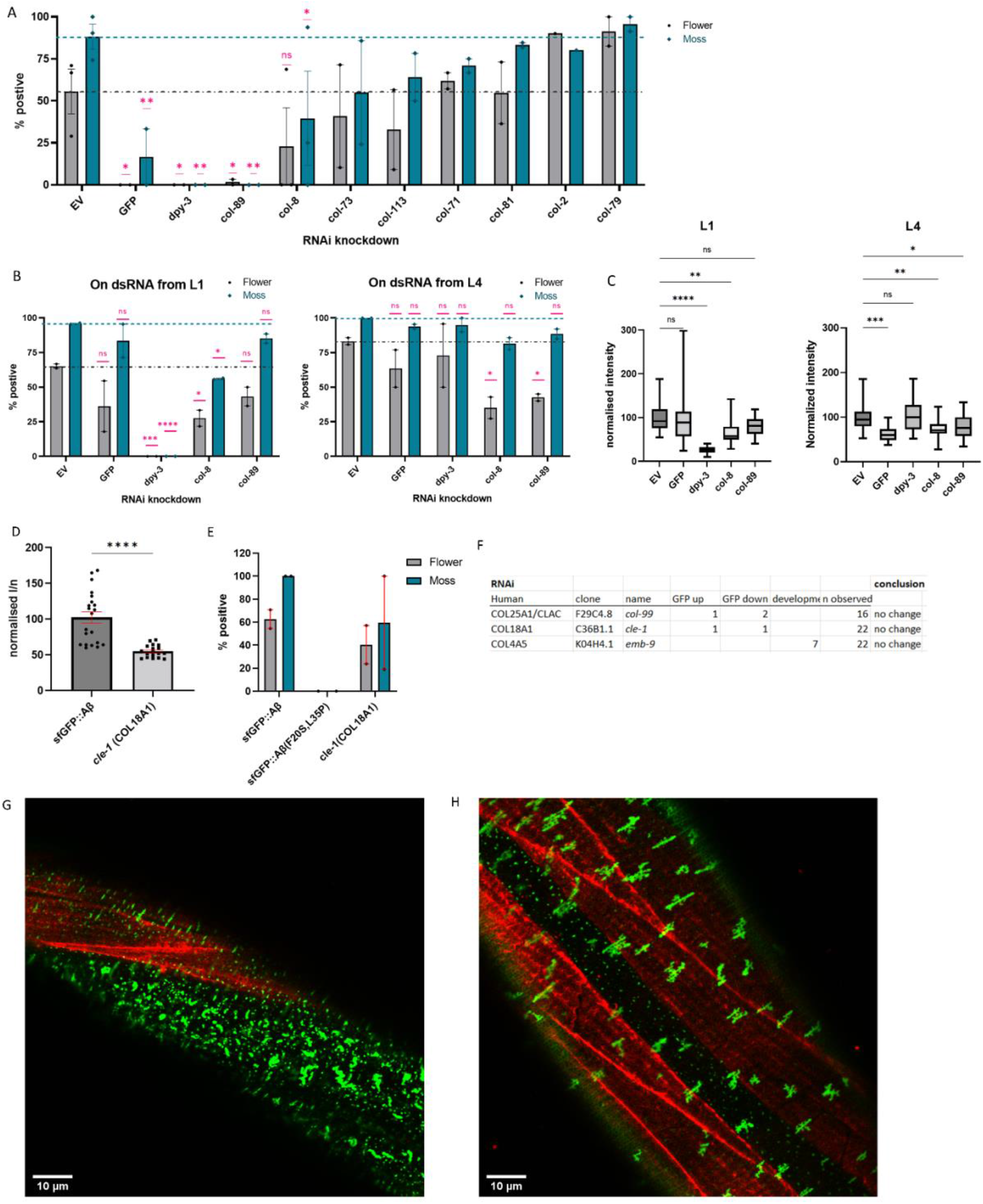
Collagens knockdown prevented or promoted extracellular sfGFP::Aβ aggregation. **A)** Collagen knockdowns that initially showed an increase in GFP intensity were followed up by observation of aggregate formation on the cuticle. RNAi of collagens *dpy-3* and *col-89* showed no sfGFP::Aβ aggregates. Statistics: ordinary two-way ANOVA, error bars SEM. **B, C)** The lack of sfGFP::Aβ aggregates could be due to a structural requirement or indirectly due to increased turnover of collagens at the ECM. To separate the two, RNAi initiated from the first larval stage (L1) was compared to RNAi initiated from the last (L4) larval stage; the latter should only take effect after the cuticle has been fully formed. **B**) Score for aggregates as the % of the population. Statistics: ordinary two-way ANOVA, error bars SEM. **C**) Normalized GFP intensity. Statistics: ANOVA, plotted: Tukey. **D, E)** Knockout of *cle-1(gk364*), the ortholog of COL18A1, showed a significant reduction of sfGFP::Aβ intensity, combined with a mild reduction in flower aggregate formation. Statistics: ANOVA, error bars SEM. **F)** Numbers represent individual trials for the categorized phenotype. n observed gives the total number of times the experiment was performed. RNAi of conserved collagens showed no effect on sfGFP::Aβ. **G)** Colocalization of sfGFP::Aβ and collagen type IV/emb-9::mCherry showed the moss aggregates in the region of the hypodermis (absence of muscle tissue underneath). **H)** Colocalization of sfGFP::Aβ and collagen type IV/emb-9::mCherry showed the flower aggregates in the region of the muscle tissue.

Next, we went back to our screening hits to determine whether the four conserved collagens (collagen type IV (*let-2* and *emb-9*), collagen type XVIII (*cle-1*), and collagen type XXV (*col-99)*), which had previously been found to influence Aβ aggregation in mammals, would also have a functional role in our system. CLE-1 has been reported to be expressed in neurons and muscles and localizes predominantly around synapses and neuromuscular junctions (Ackley et al., 2001; Heljasvaara et al., 2017). Since RNAi is incompletely penetrant and less effective in neurons (Asikainen et al., 2005), we used a mutant for this neuronal-expressed collagen. We found that collagen type XVIII orthologue *cle-1(gk364*) mutants showed lower overall sfGFP::Aβ fluorescence and a mild reduction in moss and flower aggregates (Figure 4D, 4E). Since CLE-1 does not localize to the cuticle, this suggests indirect effects of *cle-1* collagen on Aβ aggregates. For collagen type XXV, *col-99* in *C. elegans*, RNAi showed no change in sfGFP::Aβ fluorescence intensity (Figure 4F). Neither *col-99(ok1204*) mutants nor COL-99 overexpression showed any consistent effect on sfGFP::Aβ fluorescence intensity (Supplementary file 3). For collagen type IV (*let-2* and *emb-9), emb-9* RNAi showed no changes in sfGFP::Aβ fluorescence intensity but showed a mild developmental delay (Figure 4F). Assessment using RNAi from the L4 stage again showed no influence on sfGFP::Aβ fluorescence intensity (Supplementary file 1). Furthermore, overexpression of EMB-9 did not noticeably change sfGFP::Aβ aggregation, nor did sfGFP::Aβ and EMB-9::mCherry colocalize (Figure 4G, 4H, Supporting figure 4.1–4.3), which could explain why we observed no effect of collagen type IV on sfGFP::Aβ aggregation in this model. In summary, although we observed that cuticular collagens could influence Aβ aggregation and Aβ aggregate removal, the previously implicated orthologues might not directly affect sfGFP::Aβ aggregation in the cuticular ECM. Nevertheless, our data points towards collagen and ECM remodeling influencing Aβ aggregation and Aβ aggregate removal.

### TIMP and MMPs regulate Aβ removal

To define a role for collagen and ECM remodeling in the development and removal of Aβ aggregates at the cuticle, we used a broad range inhibitor of metalloproteases, batimastat (BB94) (Jacobsen et al., 2010). Consistent with our RNAi screening hits on extracellular proteases, exposure to batimastat increased the sfGFP::Aβ load, represented by an increase in GFP intensity (Figure 5A, 5B), suggesting that metalloprotease activity is essential in removal.

**Figure 5.**
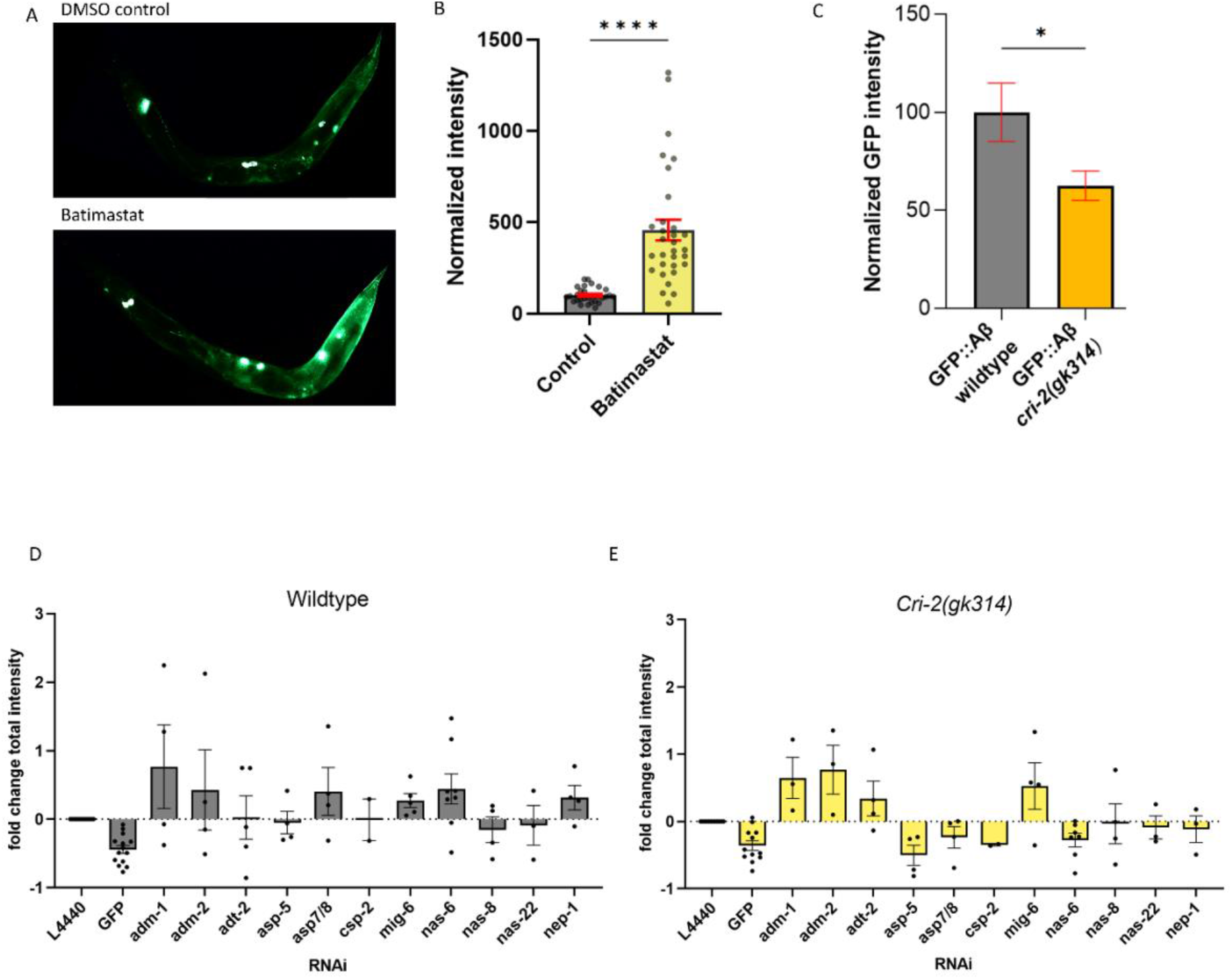
Regulation of ECM structure and turnover influences aggregates. **A, B)** Exposure to the metalloproteinase-inhibiting drug batimastat showed a marked increase in sfGFP::Aβ; suggesting reduced removal. **A**) representative image of the GFP intensity on the DMSO control treatment and representative image of the GFP intensity on the batimastat treatment **B**) Combined, normalized data of three independent trials. Statistics: unpaired *t*-test. Error bars SEM. **C)** The deletion of the Tissue Inhibitor of MetalloProteases (TIMP) *cri-2(gk314*) showed a decrease of sfGFP::Aβ load, suggesting increased activity of metalloproteases. Statistics: unpaired *t*-test. Error bars SEM. **D, E)** To determine which metalloprotease under the regulation of CRI-2 showed the most potential to assist in the removal of extracellular sfGFP::Aβ, GFP intensities per population were measured for RNAi of several individual metalloproteases and compared between the wildtype and *cri-2* mutant background. Loss of ADM-2 showed the largest increase in sfGFP::Aβ intensity and was selected for follow-up. Plotted: normalized mean of independent trials (each trial is one dot) with SEM.

Metalloprotease activity is controlled by tissue inhibitors of metalloproteases (TIMPs) (Nagase et al., 2006). Interestingly, we also picked up *cri-2* in the screen, an ortholog of human TIMP (Teuscher et al., 2019). Knockdown of *cri-2* resulted in reduced sfGFP::Aβ intensity, suggesting that when the inhibitor of metalloproteases is removed, the metalloproteases increase their activity and remove the sfGFP::Aβ. To validate this, we crossed in a genetic deletion, *cri-2(gk314*), and found lower levels of sfGFP::Aβ compared to a wild-type background at the 24h time point (Figure 5C). This reduction is not due to a difference in initial expression of sfGFP::Aβ, as there was no difference between intensities in the *cri-2* mutant and wild-type backgrounds over the first 20h after induction (Supporting figure 5.1). To identify the metalloproteases inhibited by CRI-2, we tested all MMPs identified in our screen in the *cri-2* mutant background (Figure 5D, 5E). From the nine metalloproteases tested, knockdown of *adm-1, adm-2, adt-2*, and *mig-6* resulted in increased GFP intensity in both the wild-type and *cri-2* mutant backgrounds (Figure 5D, 5E). This suggests that when the inhibitor CRI-2 is absent, these four metalloproteases become more active and contribute to the removal of sfGFP::Aβ. Knockdown of *adm-2* outperformed the other metalloproteases as indicated by the higher intensity in the *cri-2* mutant background (Figure 5E), suggesting a more prominent role for ADM-2.

### ADM-2 reduces ss::sfGFP::Aβ intensity

ADM-2 is a disintegrin plus metalloprotease family member, a membrane-bound metalloprotease with extracellular peptidase M12B, disintegrin, and EGF-like domains. ADM-2 is an ortholog of the human ADAM9, which is implicated in inflammation, cancer, and Alzheimer’s disease by cleaving the amyloid precursor protein (APP) (Chou et al., 2020), but whether ADAM9 plays a potential role in Aβ removal is unknown. To verify the increase of sfGFP::Aβ upon *adm-2* knockdown, we crossed a deleterious mutant for *adm-2(ok3178*) into the wildtype sfGFP::Aβ strain as well as into the *cri-2(gk314*) mutant strain. We confirmed that *cri-2* mutants showed lower, whereas *adm-2* mutants showed higher sfGFP::Aβ fluorescent levels (Figure 6A). The double mutants of *cri-2; adm-2* showed wild-type sfGFP::Aβ fluorescent levels (Figure 6A), suggesting that the benefits of losing the inhibitor of metalloprotease CRI-2 on Aβ aggregation are dependent on ADM-2.

**Figure 6.**
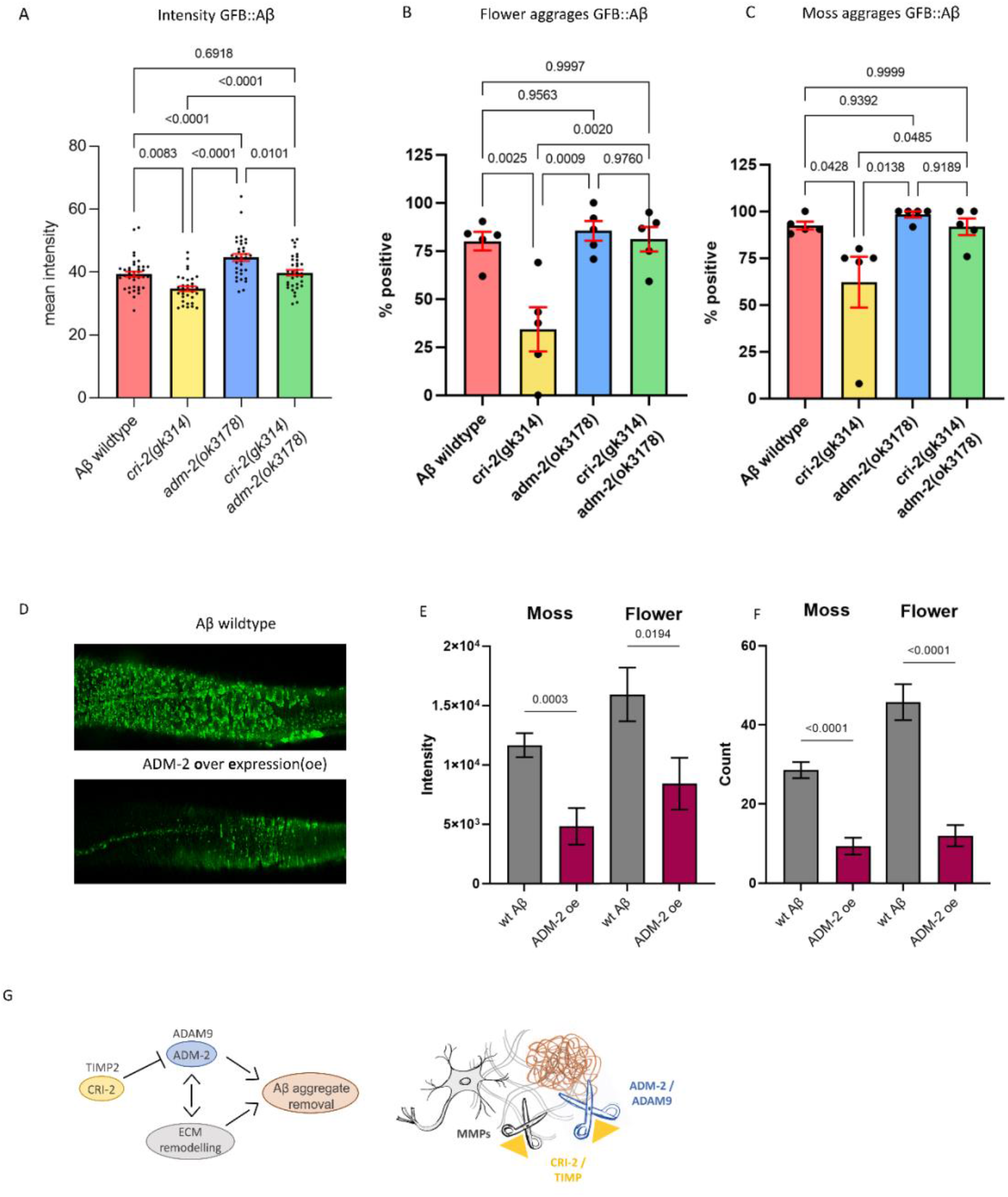
ADM-2 was required and sufficient to remove extracellular sfGFP::Aβ. **A)** The GFP intensity for sfGFP::Aβ was reduced in the *cri-2* mutant background. Loss of *adm-2* increased sfGFP::Aβ intensity, even in the *cri-2/adm-2* double mutant. Data were combined from four independent experiments. Statistics: ordinary two-way ANOVA. Plotted bars are mean with SEM. **B, C)** Similarly, loss of *cri-2* resulted in fewer animals with flower and moss aggregations, while loss of *adm-2* increased both aggregation types. Loss of *cri-2* and *adm-2* in the double mutant background showed that other metalloproteases could not compensate for the loss of *adm-2* regarding the removal of extracellular aggregates; suggesting that *adm-2* was required for the removal of extracellular sfGFP::Aβ aggregates. Statistics: ANOVA. Plotted is the percentage of the population positive for aggregation type, one dot per experiment, bars are mean with SEM. **D)** Visual representation of the observation that overexpression of *adm-2* (ADM-2 oe) led to a reduction of extracellular sfGFP::Aβ aggregates. **E, F)** Overexpression of ADM-2 was sufficient to lead to a significant reduction of sfGFP::Aβ aggregates, both in count and intensity of aggregates, measured 48h after induction of transgene expression. Data is the intensity and count measures over two independent experiments. Plotted are mean and SEM. Statistical analysis: unpaired *t*-test. **G)** Schematic representation of the mechanisms in which ADM-2, under regulation by CRI-2, either directly or indirectly assisted in removing extracellular sfGFP::Aβ aggregates.

### ADM-2 is required for Aβ aggregate removal

Next, we assessed whether the changes in GFP intensity reflect changes in secreted sfGFP::Aβ aggregates. In wild-type sfGFP::Aβ populations, about 81% showed the ‘flower’ structures (Figure 6B), and about 93% showed the ‘moss’ structures (Figure 6C). Similar to the total sfGFP::Aβ fluorescent levels at the 24 hour time point, we found that *cri-2* mutant had lower, whereas *adm-2* mutants had higher moss and flower-positive animals, which in *cri-2; adm-2* double mutants were returned to wild-type levels (Figure 6B, 6C). This confirms that the observed changes in GFP intensity reflect changes in aggregation and highlights that *adm-2* was required for the clearance of extracellular Aβ aggregates in the ECM.

### ADM-2 is sufficient for Aβ aggregate removal

To test whether ADM-2 is sufficient to reduce Aβ aggregation, we constructed a transgene with ADM-2::mScarlet-I that inducibly overexpressed ADM-2 under the control of the *hsp-16.2* promoter. The mScarlet-I fluorophore is situated at the cytoplasmic domain so as not to obstruct ADM-2 activity in the extracellular space. While the initial localization was the same as they are both induced by the same promoter, ADM-2::mScarlet-I did not colocalize with sfGFP::Aβ aggregates at 24h after induction (Supporting figure 6.1). This could be either due to the mScarlet tag being intracellular, *i.e*., only membrane-bound ADM-2 was visible, or ADM-2 efficiently removed nearby sfGFP::Aβ. ADM-2 overexpression showed a trend toward a reduced count and intensity of both moss and flower aggregates at 24h (Supporting figure 6.2). However, when the aggregate count, size, and intensity were compared 48h after induction, we found a significant reduction of all aggregation structures upon ADM-2 overexpression (Figure 6D-F). These data support the idea that ADM-2 is both required and sufficient to reduce sfGFP::Aβ aggregates in the cuticle. Thus, we propose the model that TIMP(CRI-2) inhibits ADAM (ADM-2) to either directly reduce Aβ aggregation or indirectly via ECM remodeling (Figure 6G).

## Discussion

Amyloid-beta plaques are a hallmark of Alzheimer’s Disease. Collagen is consistently found within these plaques, but a functional relation between amyloid-beta and collagens has not been shown *in vivo*. We assessed the interaction between ECM components and regulators of ECM turnover for a potential mediating role in amyloid pathology, using novel *C. elegans* transgenic strains to express and secrete human amyloid beta; Aβ_1-42_. This amyloid beta was found to form two types of extracellular aggregates associated with underlying tissue type. Targeted genetic knockdown showed mediating effects on aggregation by several extracellular proteins, including collagens and metalloproteases. A complete absence of aggregation was observed for the knockdown of *dpy-3* and *col-8* collagens. From a selection of metalloproteases, <u>A</u> <u>D</u>isintegrin and <u>M</u>etalloprotease 2 (ADM-2) were found to be most effective in reducing amyloid-beta aggregation. We found overexpression of ADM-2 to be sufficient to remove extracellular Aβ aggregates *in vivo*. These findings support a potential active, mediating role for ECM components on Aβ aggregation.

Generally, metalloproteases are known for their function in mediating ECM remodeling by cleaving collagens. While some collagens are long-lived ECM components, ECM turnover allows for damage repair (wound healing) or in response to other tissue demands, such as exercise (Ewald, 2019; Kritikaki et al., 2021; Xue & Jackson, 2015). In combination with the collagens we found, this could imply that ADM-2 assists in the removal of Aβ aggregates by remodeling the ECM. The collagen components that, when the expression is knocked down, suppress aggregate formation suggest that there are ECM composition requirements for aggregates to form. It is unclear if these specific collagens are required for direct interaction with Aβ before aggregate formation or if a more general structural composition within the ECM is required. However, these data support the concept that ECM dynamics are fundamental to Aβ aggregation, and ECM remodeling contributes to aggregate removal.

Another category identified in the RNAi screen bridges the extracellular and intracellular environments and involves clathrin-mediated endocytosis, the sorting endosome, and (targeted) vesicle secretion. In our screen, knockdown of RAB-7 led to an abundant accumulation of small, bright green vesicles near the cuticle, while aggregates remained absent (Supporting figure 3). Furthermore, members of the ESCRT complex, as well as SNAREs and sorting nexins, were indicated to alter the localization, accumulation, and aggregate formation of Aβ. Conceptually, ECM remodeling and vesicle uptake and secretion may serve a common purpose regarding dynamic ECM adaptations. In the process of ECM turnover, metalloproteases are actively cleaving ECM components such as collagens and fibronectins (Shi & Sottile, 2011), and the resulting cleaved products are internalized via receptor-mediated phagocytosis and degraded in the lysosome (Arora et al., 2000). Furthermore, to secrete metalloproteases, newly synthesized collagens, or Aβ to the ECM, the sorting endosome and (vesicle) secretion pathways are in play (Chang et al., 2021). As such, these seemingly distinct mechanisms could all work together for the collective purpose of extracellular aggregate removal.

In recent work, ADM-2 overexpression was shown to lead to molting defects (Joseph et al., 2021). To allow growth, the cuticle of *C. elegans* is shed and replaced by a new cuticle, secreted and deposited by hypodermal cells underneath the cuticle. Initiation of the molt requires the internalization of sterol hormones and activating a cascade of proteases to mediate the shedding of the old cuticle. As such, molting depends on ECM remodeling, in which ADM-2 plays an essential role. Interestingly, one of the potential targets of ADM-2 cleavage revealed in that work is LRP-1, the *C. elegans* low-density lipoprotein receptor orthologous to human LRP1 (Joseph et al., 2021). LRP-1 in *C. elegans* is a membrane-bound receptor, which can sequester sterols from the extracellular environment, and when internalized together, these sterols can initiate molting (Yochem et al., 1999).

ADM-2 is suggested to cleave LRP-1 and release it from the membrane, then referred to as sLRP. Although this sLRP-1 can still capture sterols, they are not internalized, leading to incomplete shedding of the cuticle (Joseph et al., 2021). Genes involved in *C. elegans* molting that are a hit in our screen are *dab-1, hgrs-1*, and *apl-1*. DAB-1 is a cytoplasmic adaptor protein involved in endocytosis. Endocytosis of sterols is essential for molting (Lažetić & Fay, 2017). HGRS-1 is a Vps27 ortholog, which recruits ESCRT machinery to endosomes. Inhibition of HGRS-1 leads to molting defects (Lažetić & Fay, 2017). Interestingly, HGRS-1 and ADM-2 colocalize (Joseph et al., 2021), which suggests they may be involved in similar pathways through direct interaction. Loss of APL-1, the APP ortholog, causes lethal defects upon shedding the cuticle, which is rescued by the expression of the extracellular part of APL-1 (Hornsten et al., 2007). The association of multiple genes involved in the molting process with an increase in sfGFP::Aβ load and aggregate formation in adult *C. elegans* suggests that changes in ECM dynamics can influence amyloid aggregate load. However, the exact role of ADM-2 and its potential targets needs further refining.

The human ortholog of ADM-2, ADAM9, has been implicated in AD and is suggested to regulate the shedding of APP as an alpha-secretase, either indirectly by regulating ADAM10 or by functioning as an alpha-secretase itself, cleaving APP in a non-amyloidogenic manner (Asai et al., 2003; Moss et al., 2011). The cleavage site for alpha-secretase is situated in the middle of the Aβ fragment, potentially allowing direct cleavage of Aβ peptides. Moreover, human ADAM9 can be alternatively spliced, losing its transmembrane and cellular domains, resulting in an extracellular, active enzyme (Hotoda et al., 2002). This could potentially be a way for ADM-2 to reach the Aβ aggregates in the cuticle. As an ortholog of ADAM9, ADM-2 could potentially cleave Aβ directly and, as such, assist in the removal of extracellular aggregates.

In conclusion, we established an *in-vivo* model to trace Aβ aggregation in the extracellular matrix. Our findings suggest that activating ECM remodeling promotes aggregate removal and could become an important strategy to ameliorate AD disease progression.

## Materials and Methods

### Strain handling

Preparation of NGM agar plates, feeding with OP50 *Escherichia coli*, and handling of *C. elegans* strains by picking as described by Stiernagle (Stiernagle, 2006). For maintenance of *C. elegans* strains, ten adults are picked and transferred to a fresh plate per generation and are kept at 15°C.

### *C. elegans* strains

For the generation of the strain expressing the transgene sfGFP::Aβ, the germline of *C. elegans* N2 Bristol (wild type) was injected with the plasmid pLSD134 at 50 ng/μL and pRF4 *rol-6(su1006*), also at 50 ng/μL. The total concentration of DNA in the injection mix was 100 ng/μL. Plasmid pLSD134 was cloned by VectorBuilder. Detailed plasmid sequence and map are in Supplementary file 4. The extrachromosomal array was integrated into the genome using UV irradiation with the Stratagene UV Stratalinker 2400 (254 nm). The resulting integrated strain was backcrossed with N2 Bristol four times and named LSD2104, which was used throughout this study.

To attain deletion of the genes *cri-2, adm-2, col-99 and cle-1* in LSD2104(sfGFP::Aβ), LSD2104 was crossed to VC718 *cri-2(gk314) V*, RB2342 *adm-2(ok3178) X*, RB1165 *col-99(ok1204) IV*, and VC855 *cle-1(gk364) I;* resulting in the strains LSD2165, LSD2201, LSD1056, and LSD1052 respectively. The double mutant background with deletions for *cri-2(gk314*) and *adm-2(ok3178*) was generated by crossing LSD2165(sfGFP::Aβ, *cri-2(gk314) V*) with RB2342 *adm-2(ok3178) X*, resulting in the strain LSD2204.

To induce overexpression of ADM-2 in LSD2104(sfGFP::Aβ) background, LSD2104 was injected with the plasmid pLSD170, *hsp-16.2p*::adm-2::mScarlet-I. This plasmid was designed to express ADM-2 with the mScarlet-I tag in the cellular compartment, so as not to obstruct enzymatic function. Plasmid pLSD170 was cloned by VectorBuilder, and a detailed plasmid sequence and map are in Supplementary file 4. Injection of pLSD170 was performed with a total DNA concentration of 50 ng/μL. Exposure to UV to induce integration was not successful, and a non-integrated line, not exposed to UV, was maintained by selecting for expression of ADM-2::mScarlet-I, resulting in the strain LSD3014.

More details on strains and primers for identification of strains are in Supplementary file 4.

### Induction of ss::sfGFP::Aβ expression

Age-synchronized populations, as described by Teuscher (Teuscher, Statzer, et al., 2019) were grown at 20°C on NGM plates for 4 days until the young-adult stage. The heat shock was performed by placing the plates at 33°C for two hours, after which they were returned to 20°C. The assessment was 24h after heat shock induction unless otherwise mentioned.

### Assessment of GFP intensity

The intensity of the GFP signal was obtained from imaging with an upright bright field fluorescence microscope, camera, and filter set according to (A. Teuscher & Ewald, 2018). Analysis software used is Fiji(Schindelin et al., 2012), with a program described in GitHub (Statzer et al., 2021), code accessible on github.com/JongsmaE/GreenIntensityCalculator. Briefly, the triple filter set is used to separate autofluorescence in the *C. elegans* gut from the GFP signal, and the autofluorescence appears yellow. The GFP intensity is calculated by the program as follows: the color image is split into green, blue, and red channels. Since yellow is an addition of green and red, green pixels are only counted if ‘green intensity’ > ‘red intensity’, and the red value is subtracted from the green value. The remaining green values are added up per selected area (worm) to obtain the total intensity. The number of pixels is counted as well to calculate the average intensity per pixel, in case one wants to account for animal size. A minimum of 20 animals are measured per condition.

### RNAi screen

RNA interference plates are prepared as described before with the addition of carbenicillin (50 μg/mL) and Isopropyl -D-1-thiogalactopyranoside (IPTG) (1 mM) after autoclaving. These plates are seeded with bacteria carrying RNAi. These bacteria originate from the Vidal RNAi library (Rual et al., 2004) and Ahringer RNAi library (Fraser et al., 2000; Kamath et al., 2003). Clones were copied from the Vidal and Ahringer libraries by growth overnight at 37°C on an LB-agar plate containing carbenicillin (50 μg/mL) and tetracycline (12.5 μg/mL) (carb/tet) and consequently grown in liquid LB(+carb/tet) from a single colony. Sequence-confirmed glycerol stocks were stored at −80°C. For RNAi experiments, clones are selected from the frozen library at −80°C and grown overnight in liquid LB with ampicillin 50 μg/mL and tetracycline (12.5 μg/mL). The following day, the cultures were spun down, and the LB(amp/tet) medium was refreshed and filled to 5x the original volume. The cultures are allowed to grow for 3h, aiming to harvest them in the growth phase. They are then concentrated 20x and resuspended in LB supplemented with 1mM IPTG to induce replication of the RNAi. These were then seeded onto the 6 cm RNAi plates, 500 μL each. Approximately 40 animals were allowed to feed at 20°C from larval stage 1 to young adulthood before heat-shock induction. As a negative control, the empty vector pL4440 was used. As a control for RNAi induction, a vector carrying RNAi for GFP is used. For the RNAi of genes that led to developmental delay, RNAi was repeated from the L4 stage to re-assess an effect on sfGFP::Aβ intensity. Raw data are presented in Supplementary file 1, together with the list of hits, including developmental delay, in separate tabs of the file.

### Confocal imaging and FRAP

Images of aggregation were taken using an upright confocal laser scanning microscope (CLSM) as described by Hess (Hess et al., 2019). Adaptations: 60/1.00 oil objective, excitation at 488 nm, an intensity of 0.3%, and a 2% agarose pad. Additionally, photobleaching was performed by exposing a selected region to UV (405nm) laser at 3% intensity for 17 seconds. Photobleached areas were observed for recovery of fluorescent signal from the first seconds up to four hours after photobleaching.

### Correlative light and electron microscopy (CLEM)

*C. elegans* were fixed with 4 % formaldehyde and 0.1 % glutaraldehyde in 0.1 M sodium cacodylate buffer, immersed in gelatine 12%, cryoprotected with 2.3 M sucrose. Samples were frozen in liquid nitrogen, and using an ARTOS 3D ultracut system equipped with a cryochamber EM UC7 Leica, 110 nm ultrathin cryosections were collected on 7×7 mm silicon wafers with fluorescent beads (PS-Speck, ThermoFisher). Light microscopy images were acquired with a widefield microscope, Thunder Leica, objective 100x/1.44. Electron microscopy images from the very same section were taken with a SEM Auriga 40 Zeiss microscope at an acceleration voltage of 800 eV, with an InLens detector, pixel size 4nm, and dwell time 100 us. Registration and alignment of the light and electron microscopy images were done with TrakEM2 (Cardona et al., 2012) within the open-source platform Fiji (Schindelin et al., 2012).

## Author contributions

All authors participated in analyzing and interpreting the data. CYE and EJ designed the experiments. JMM and EJ performed the CLEM. EJ performed all other experiments. EJ and CYE wrote the manuscript in consultation with JMM.

## Author Information

The authors have no competing interests to declare. Correspondence should be addressed to CYE.

## Acknowledgment

We thank Victoria Brügger for her help with screening and Charlotte Meneghin for helping with the validation of the screen hits. Some *C. elegans* strains were provided by the CGC, which is funded by the NIH Office of Research Infrastructure Programs (P40 OD010440). We thank David S. Fay for the helpful discussions. We thank Ursula Lüthi for cyrosectioning. EJ was funded for the first three years by a grant from the Mibelle Group Biochemistry, Mibelle AG, Switzerland. Funding from the Swiss National Science Foundation PP00P3_163898 and 190072 to CYE and 190072 to EJ.

---

**Supporting Figure 1.1.**
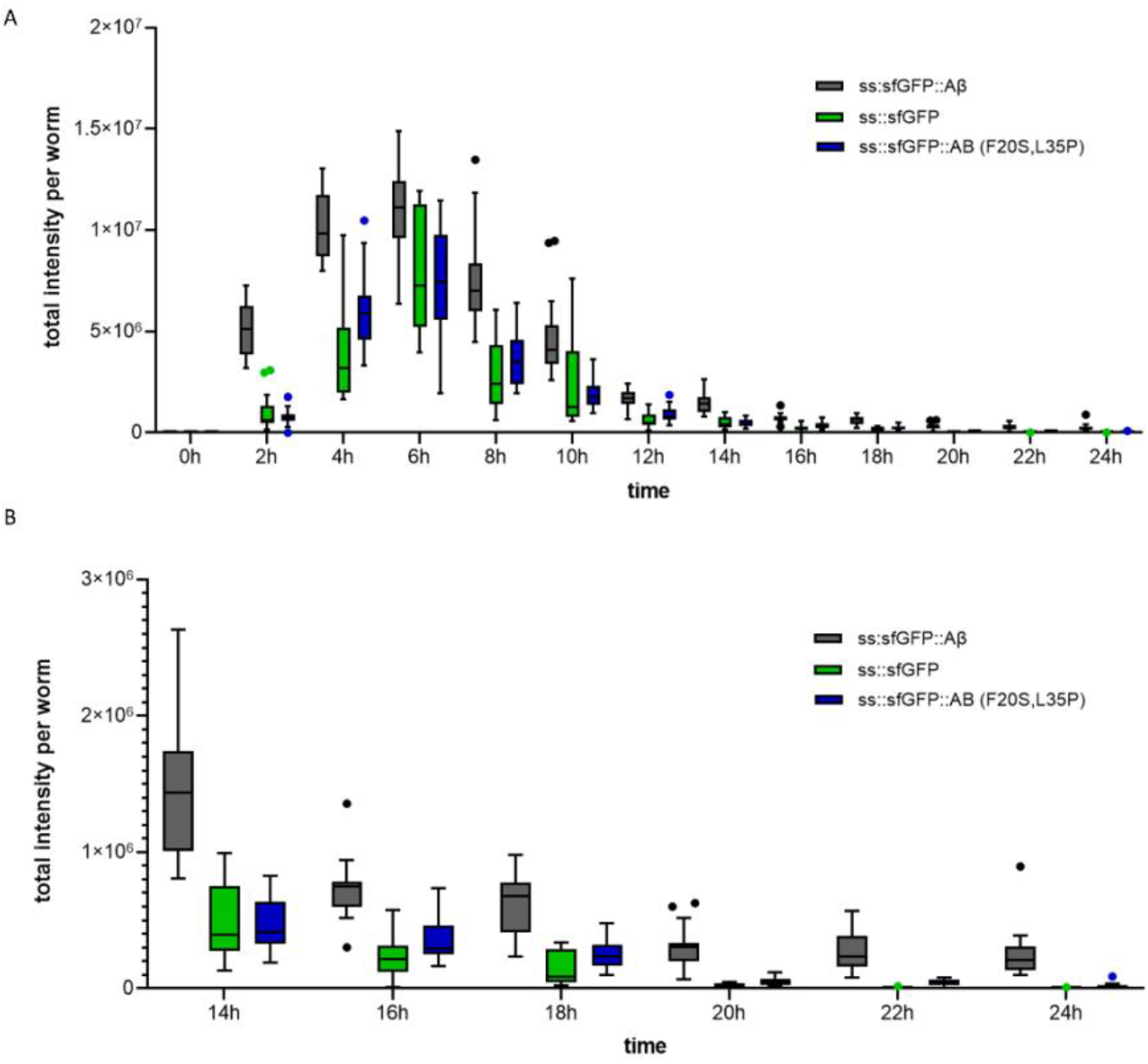
Time course of secreted amyloid beta. **A)** GFP-only and non-aggregating amyloid-beta showed similar induction as wild-type amyloid-beta. However, in contrast to wild-type amyloid-beta, GFP-only and non-aggregating amyloid-beta are efficiently removed within a 24h timespan. **B)** Tail end of A for better visibility. Plot: Tukey.

**Supporting Figure 1.2.**
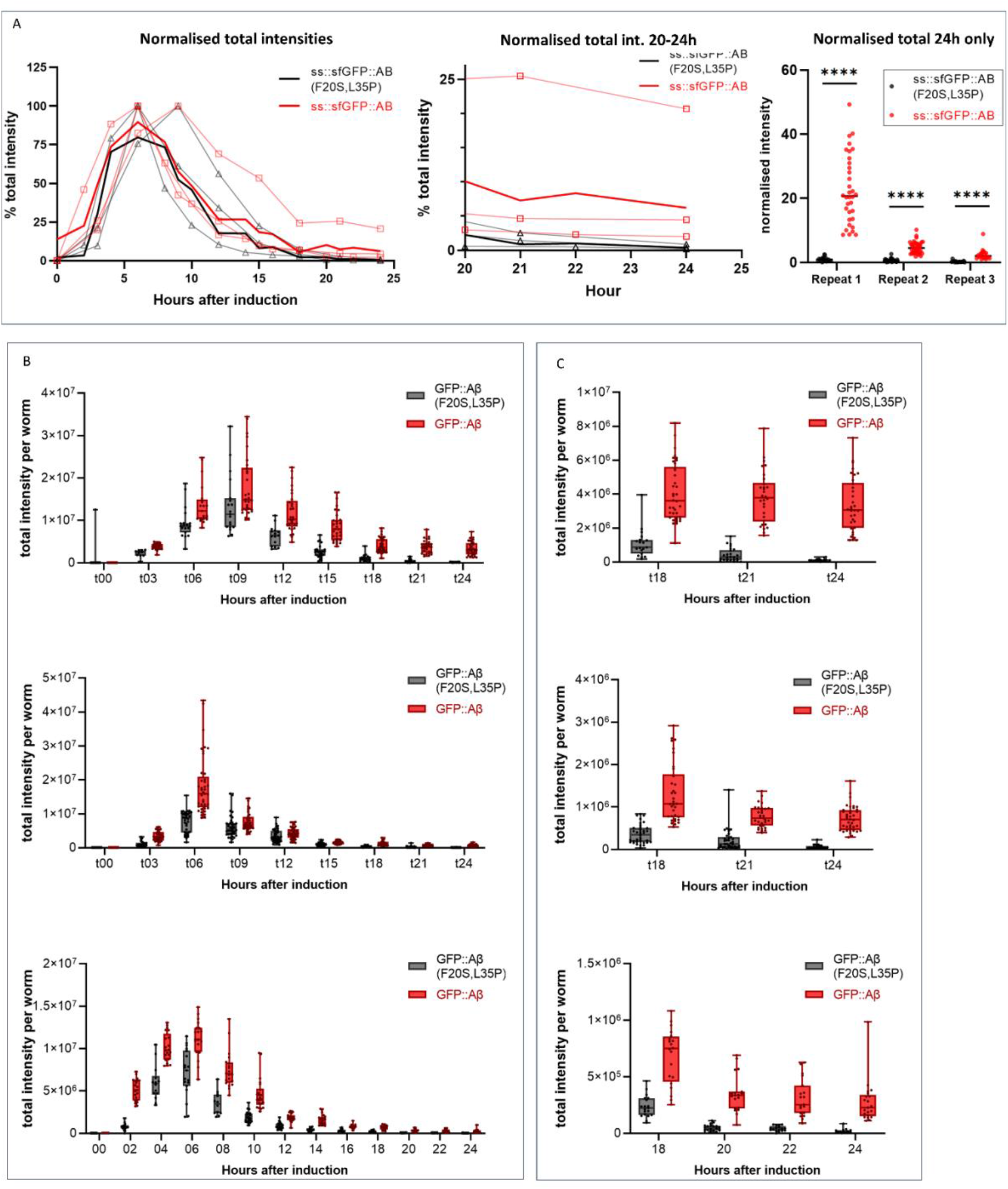
Quantification of the secreted amyloid-beta time course. Three independent trials were used to create the average for the main figure. Raw and normalized intensities are available in the data source file. **A)** Independent repeats are shown together with the average used as the main figure. Each trial was normalized to its peak (100%) intensity. Sub III contains the 24h timepoint for each repeat. The statistical test used was an unpaired, two-tailed *t*-test. ****: p < 0.0001 for each. **B)** Raw intensity per independent trial, showing the variation within each. Plot: Tukey. **C)** The last few time points for each repeat showed the sfGFP::Aβ(F20S, L35P) values are still going down, while sfGFP::Aβ seems to plateau. Plotted: Tukey.

**Supporting Figure 2.1.**
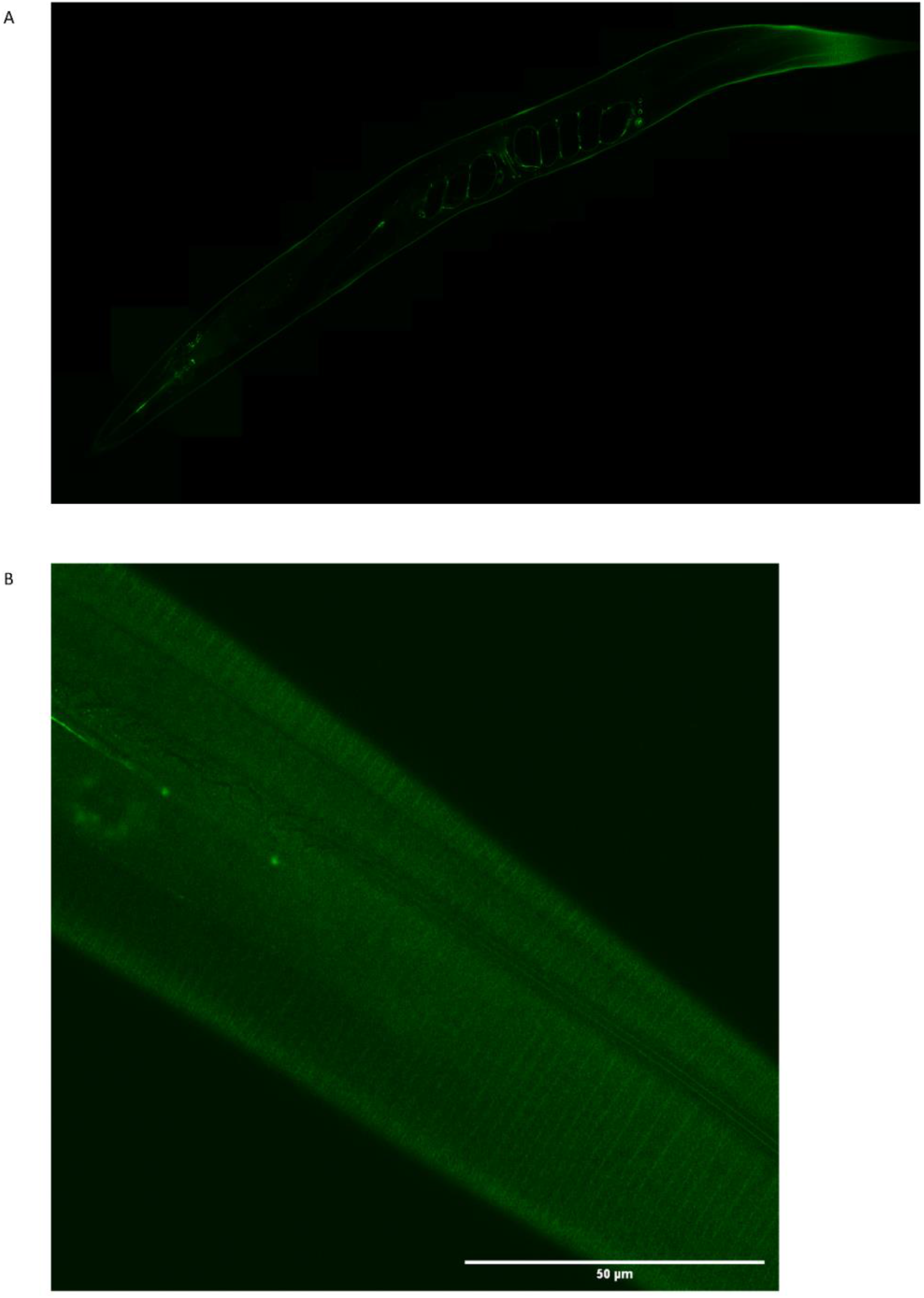
Flower and moss structures were not observed in the GFP-only strain. The GFP-only expression appears smooth and uniform in the animal and cuticle. **A)** Full-length GFP-only image, taken 16h after induction of expression. **B)** view at the cuticle of the GFP-only strain, 16h after induction, brightness enhanced 40%.

**Supporting Figure 2.2.**
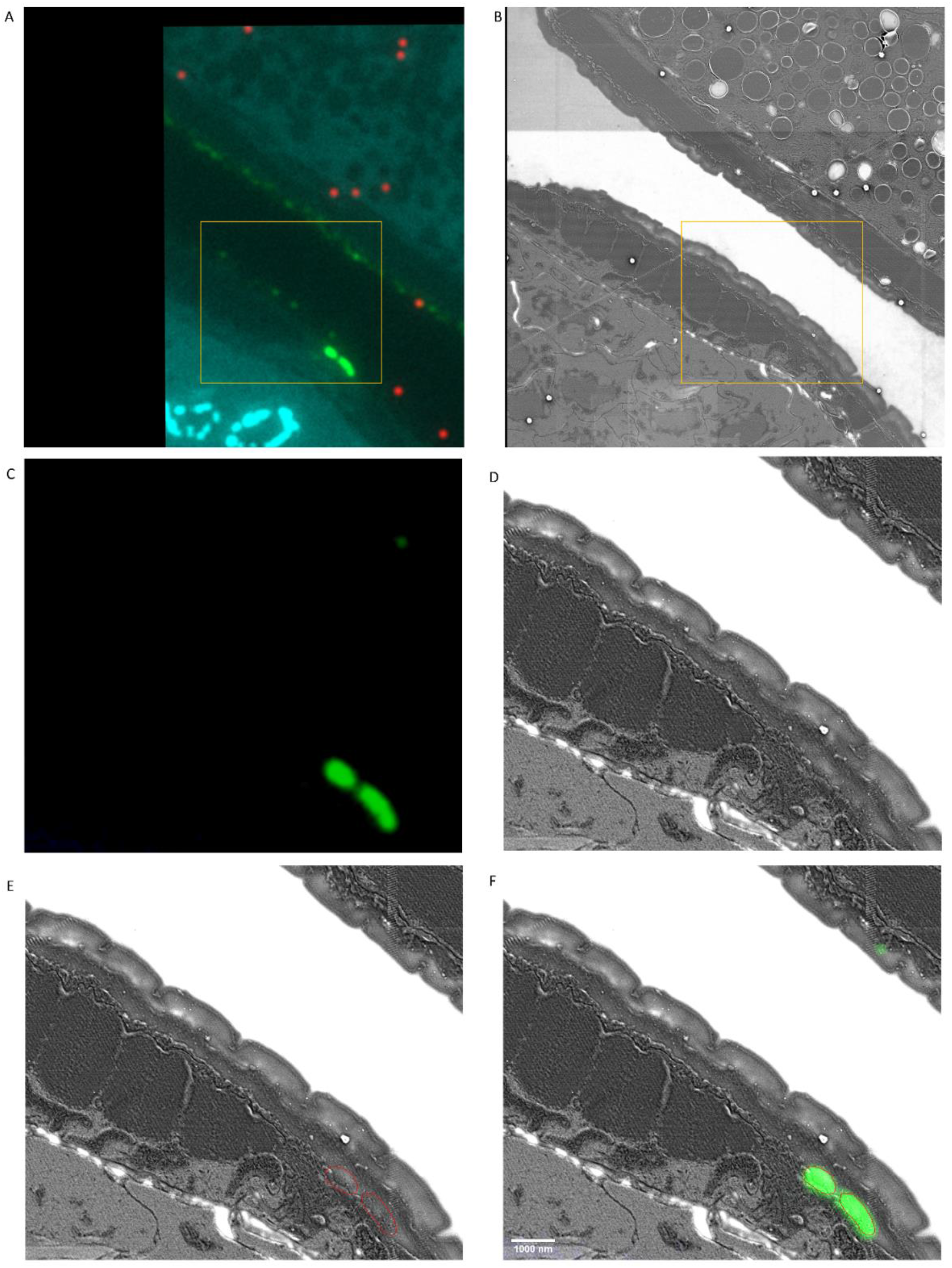
Location of sfGFP::Aβ apical to the hypodermis. **A)** Unedited fluorescence image. **B)** Unedited EM image. **C)** Crop of- and intensity reduced fluorescence image. **D)** Corresponding crop of the EM image, contrast adjusted. **E)** Overlay of EM image with the contour of GFP signal. **F)** Composite image of EM and fluorescence-adjusted, cropped images. Scale bar is 1000 nm.

**Supporting Figure 2.3.**
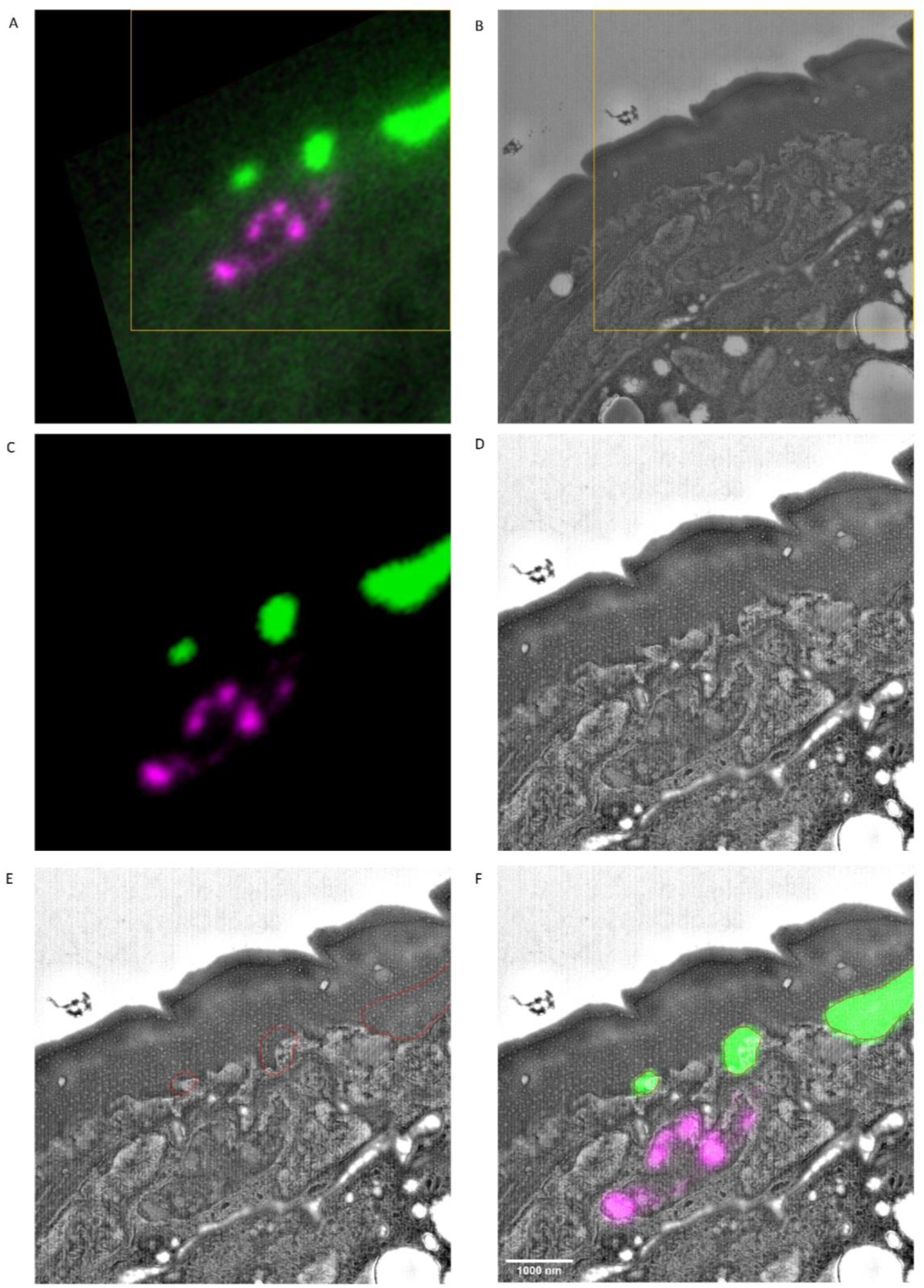
Location of sfGFP::Aβ above the hypodermis. **A)** Unedited fluorescence image. **B)** Unedited EM image. **C)** Crop of- and intensity reduced fluorescence image. **D)** Corresponding crop of the EM image, contrast adjusted. **E)** Overlay of EM image with the contour of GFP signal. **F)** Composite image of EM and fluorescence-adjusted, cropped images. Scale bar is 1000 nm.

**Supporting Figure 2.4.**
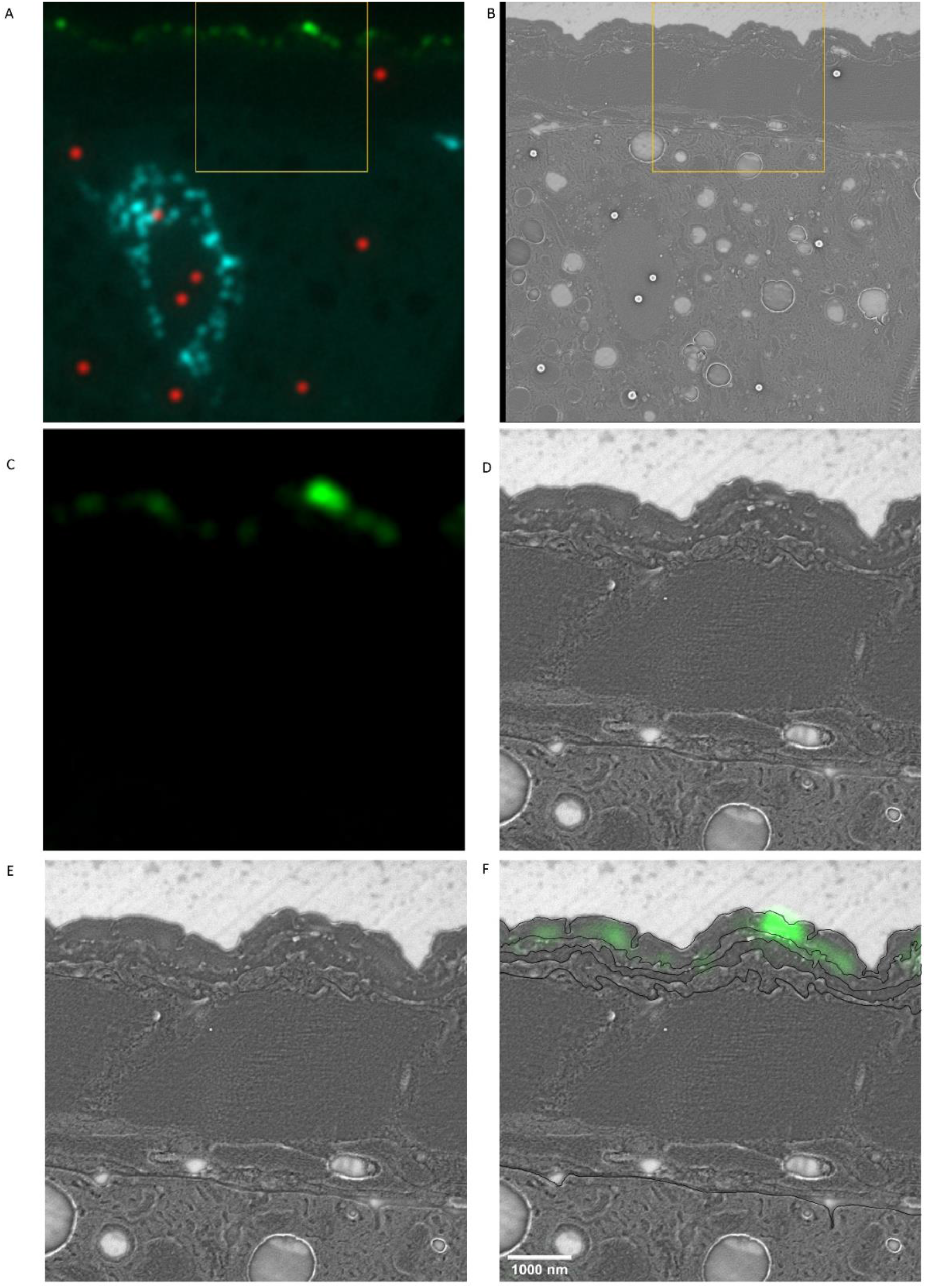
Location of sfGFP::Aβ in the cuticle. **A)** Unedited fluorescence image. **B)** Unedited EM image. **C)** Crop of- and intensity reduced fluorescence image. **D)** Corresponding crop of the EM image, contrast adjusted. **E)** Overlay of EM image with the contour of GFP signal. **F)** Composite image of EM and fluorescence-adjusted, cropped images. Scale bar is 1000 nm.

**Supporting Figure 3.1.**
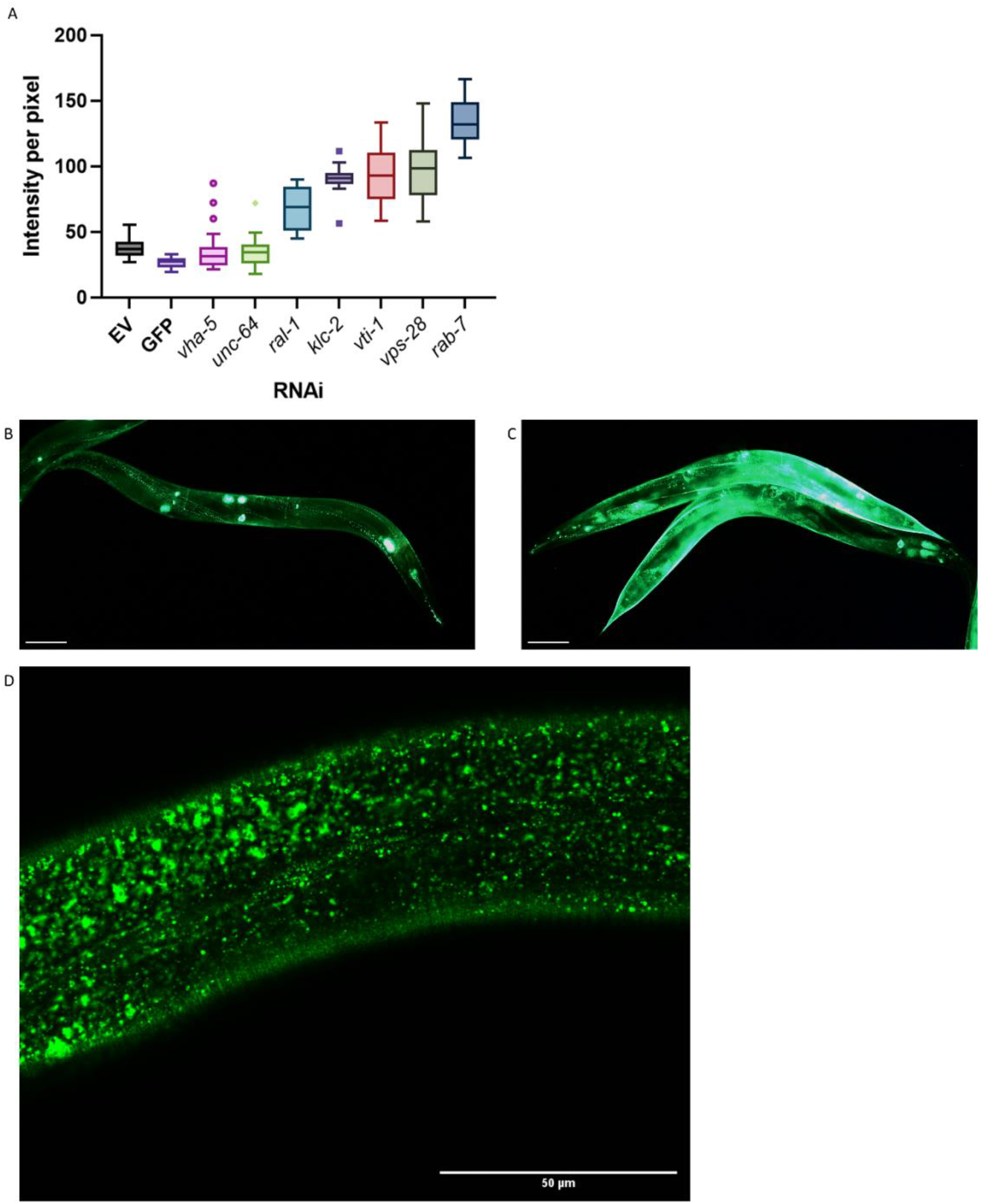
Vesicle and transport screening hits might prevent extracellular amyloid beta aggregation simply by blocking export. **A)** Intensity per pixel for all follow-up hits for the ‘vesicle’ group; genes involved in transport, secretion, or uptake of vesicle pathways. Similar results for the genes shown in A but here, as a representative, the follow-up results are only shown for *rab-7* knockdown. Knockdown by *rab-7* RNAi showed high intensity of the GFP signal, yet no flower or moss aggregates at the cuticle. **B, C)** RNAi of *rab-7* (representative image C) showed a remarkable increase in GFP intensity compared to the empty vector (representative image B). Scale bars are 50 μm. **D)** *rab-7* RNAi showed a high density of vesicles near the cuticle, yet no moss or flower pattern of aggregation. Scale bar is 50 μm.

**Supporting Figure 4.1.**
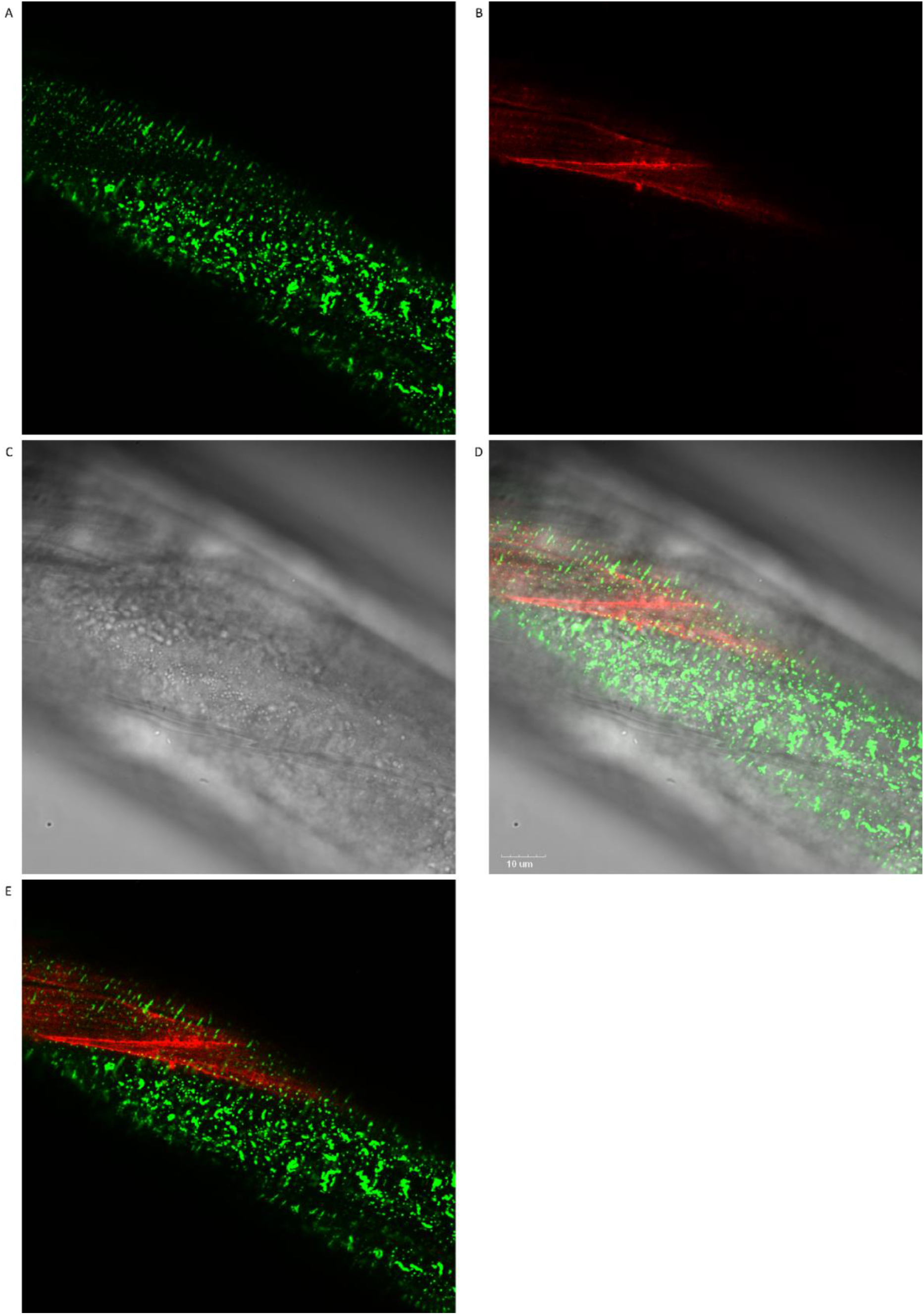
Co-localization of sfGFP::Aβ and collagen type IV/emb-9::mCherry revealed that the moss aggregates consistently colocalize to the hypodermis in the absence of muscle tissue underneath. **A)** GFP signal image. **B)** mCherry signal image. **C)** Normal light image. **D)** All three merged images. Scale bar 10 μm **E)** GFP and mCherry signals fused.

**Supporting Figure 4.2.**
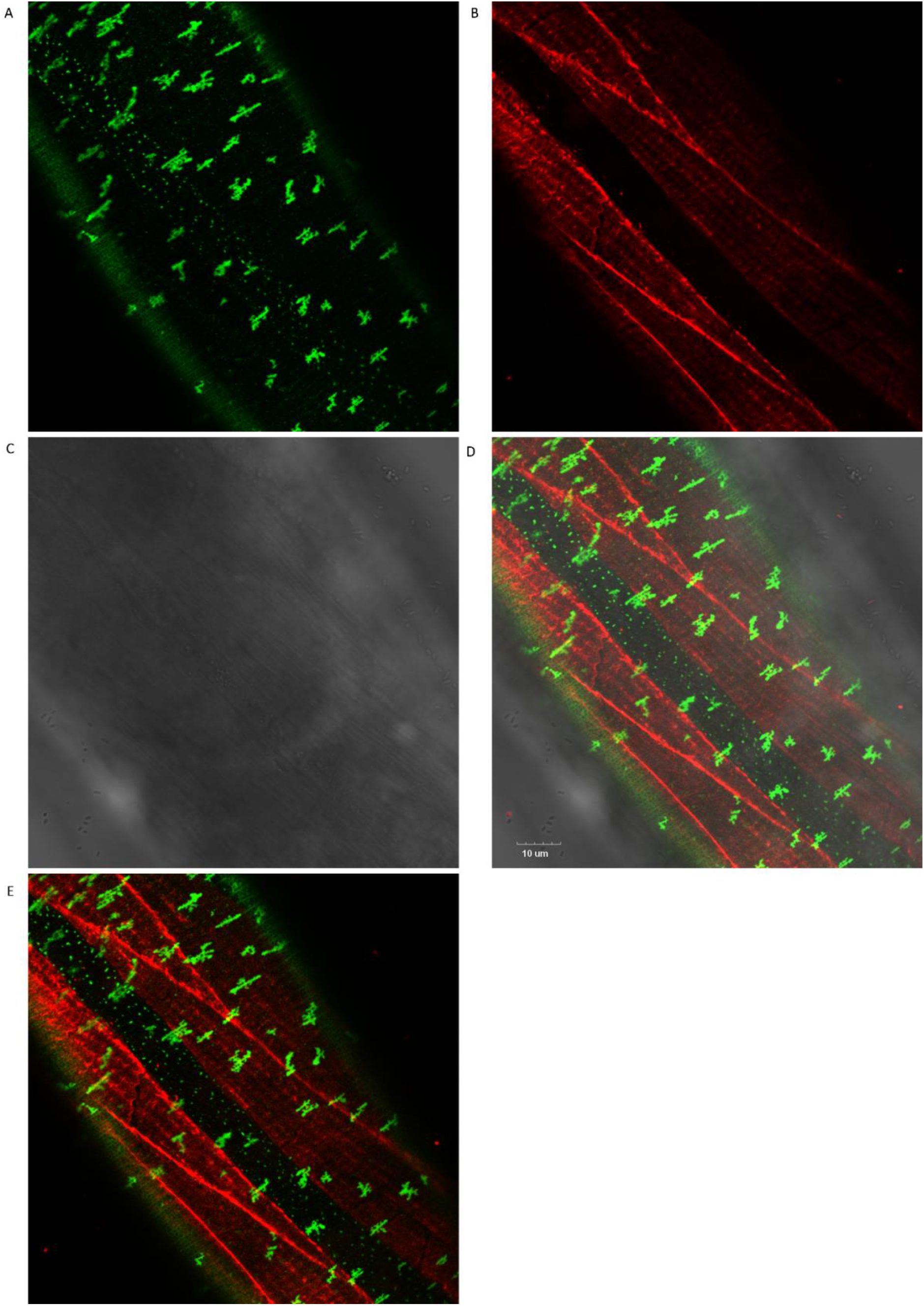
Colocalization of sfGFP::Aβ and collagen type IV/emb-9::mCherry revealed the flower aggregates consistently colocalize above the muscle tissue. **A)** GFP signal image, **B)** mCherry signal image **C)** Normal light image **D)** All three merged images, scale bar 10 μm **E)** GFP and mCherry signals fused.

**Supporting Figure 4.3.**
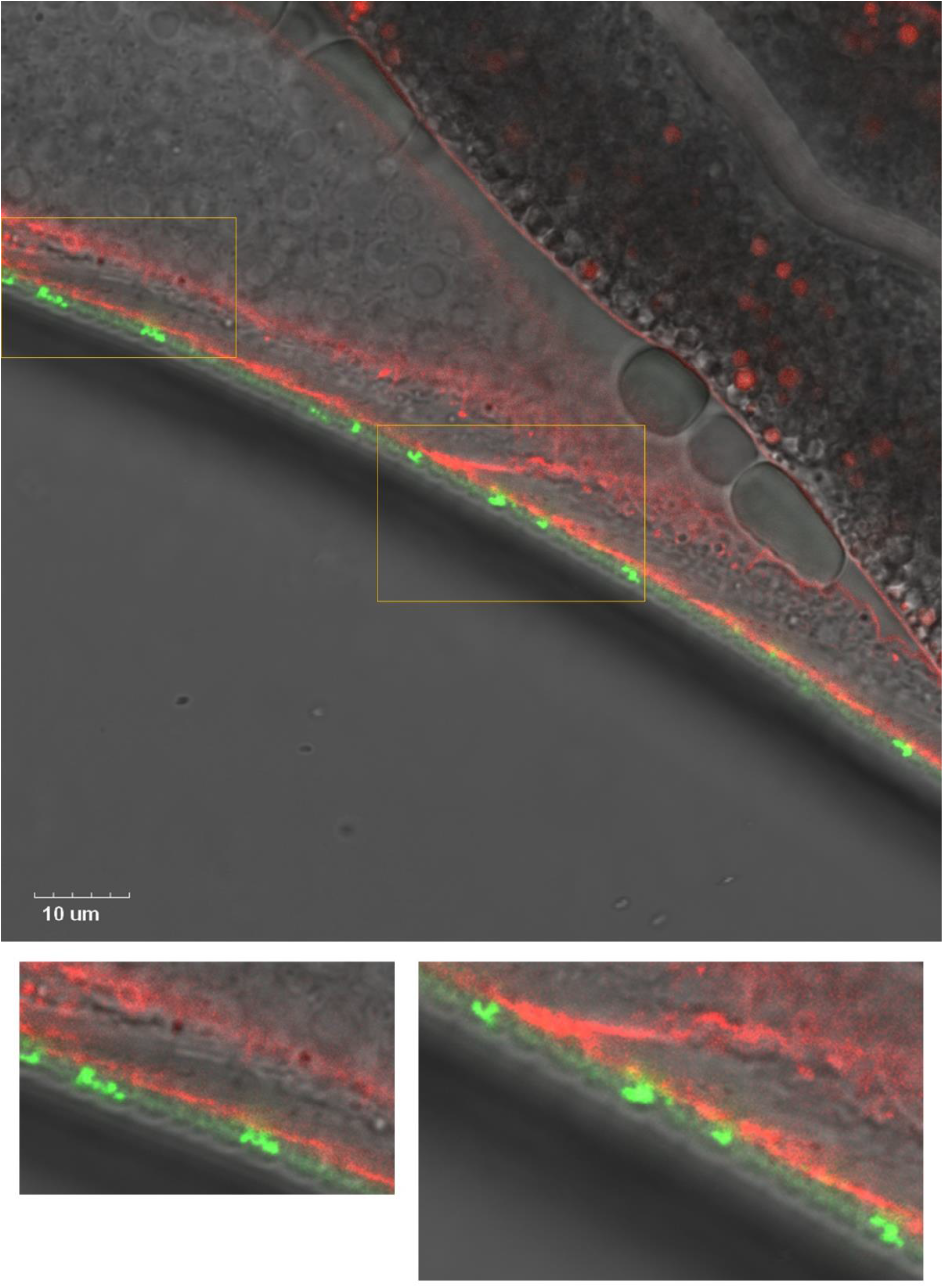
Colocalization of sfGFP::Aβ and collagen type IV/emb-9::mCherry showed that they do not colocalize. Rather, the GFP seems to localize to the cuticle while the mCherry is situated just below the cuticle. Scale bar 10 μm.

**Supporting Figure 5.1.**
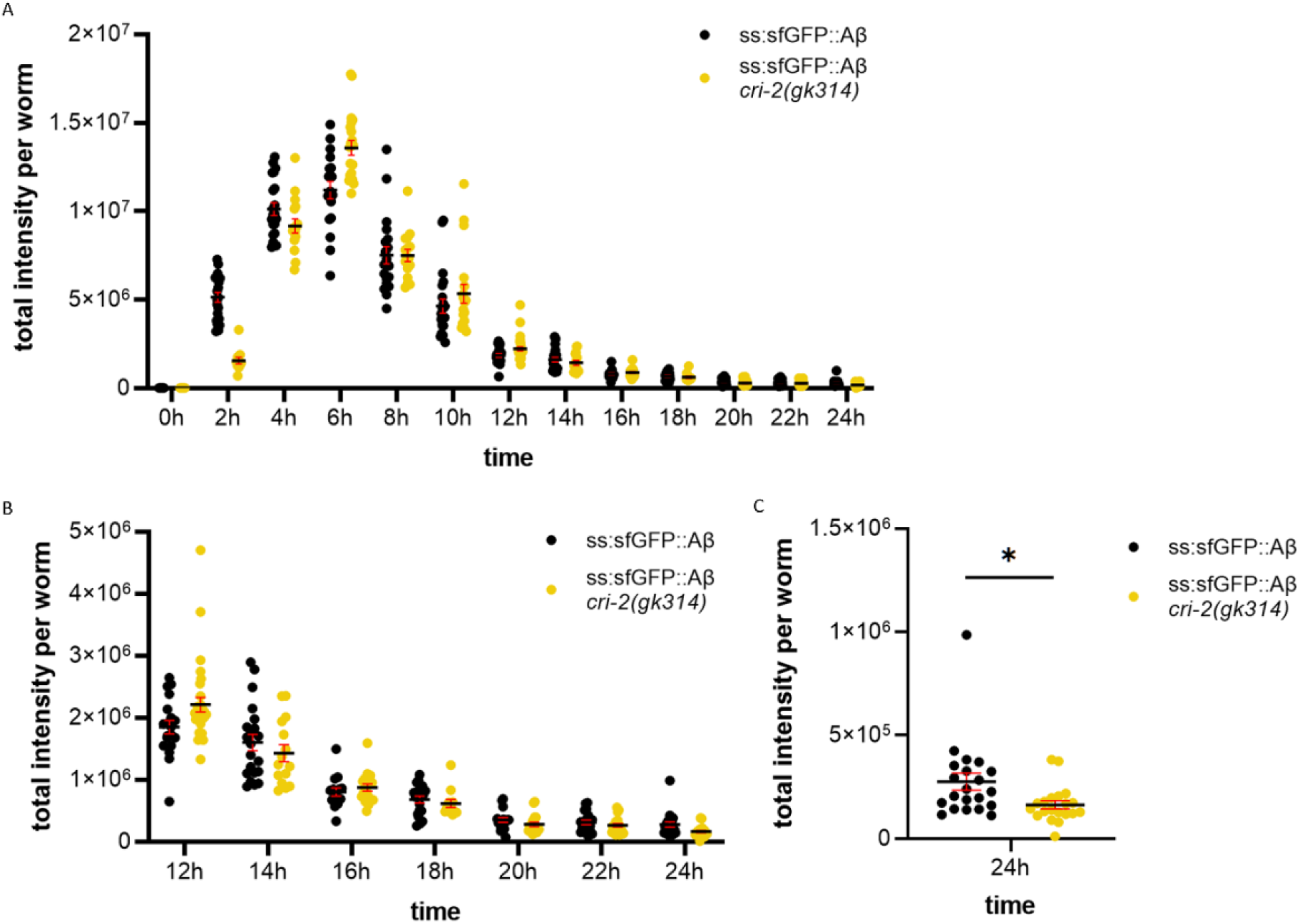
Secreted sfGFP::Aβ intensities were similar in induction between wildtype and *cri-2(gk314*) backgrounds, yet, at 24h, less GFP signal is found for the *cri-2(gk314*) background. Black line: mean. Error: SEM. **A)** total intensity measured per worm every second hour after induction up to 24h. Every point represents the GFP intensity of one worm. Black line: mean. Error: SEM. **B)** Cropped 12-24h for better resolution of the tail end of the graph in A. **C)** cropped to the 24h timepoint for better resolution of the difference between wildtype and *cri-2* mutant background. Statistics: unpaired *t*-test, *P* value 0.0201.

**Supporting Figure 6.1.**
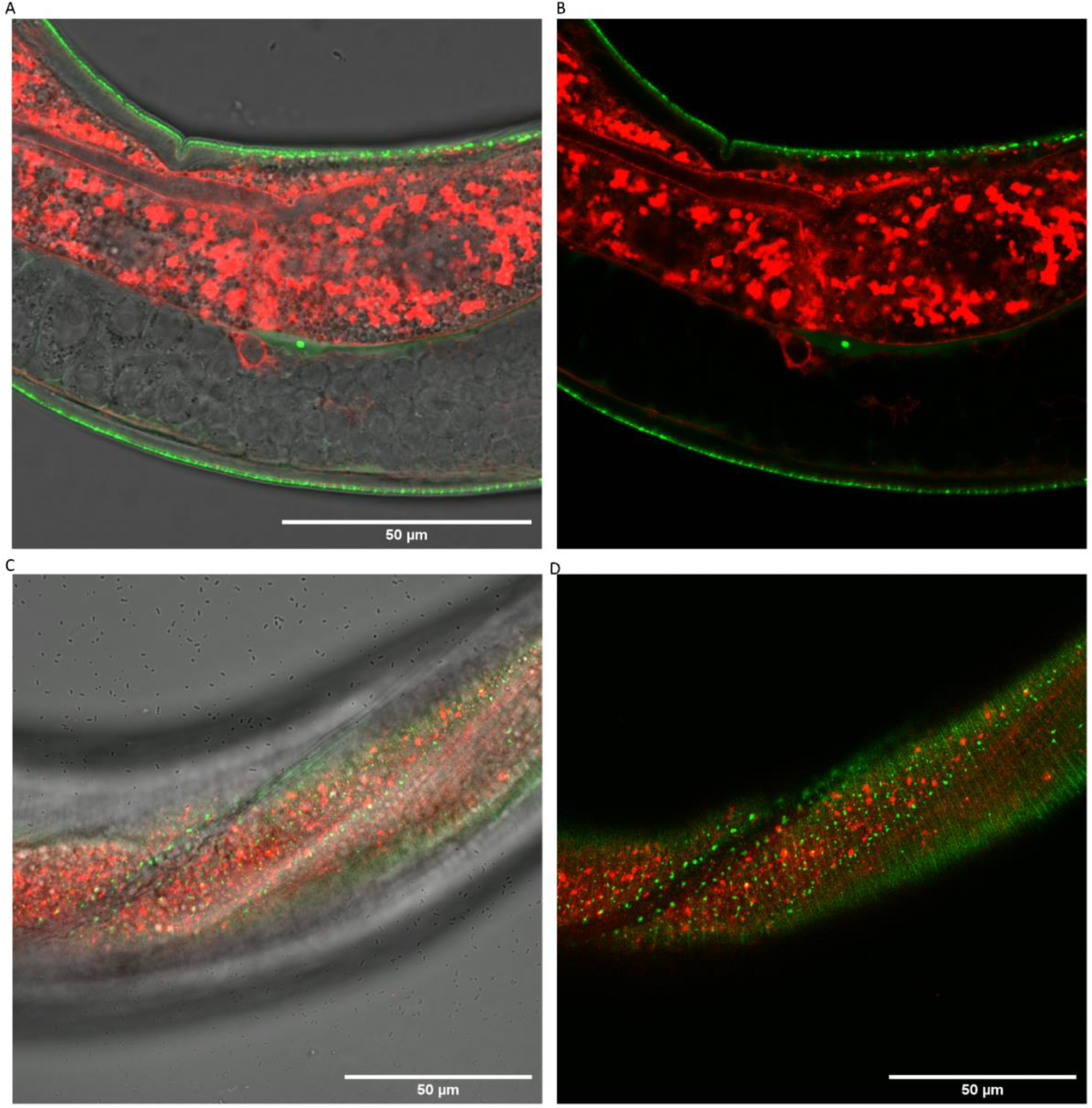
Intracellular ADM-2::mScarlet-I did not colocalize with sfGFP::Aβ. **A, B)** No colocalization of intracellular ADM-2::mScarlet-I and secreted sfGFP::Aβ aggregates **A)** Merged image of normal light, GFP and mScarlet-I **B)** The same image as A, without the normal light, merged mScarlet-I and GFP signal. **C, D)** No colocalization of ADM-2::mScarlet-I and sfGFP::Aβ was observed at the cuticle. **C)** Merged image of normal light, GFP, and mScarlet-I **D)** The same image as C, without the normal light, merged mScarlet-I and GFP signal. Scale bars 50 μm.

**Supporting Figure 6.2.**
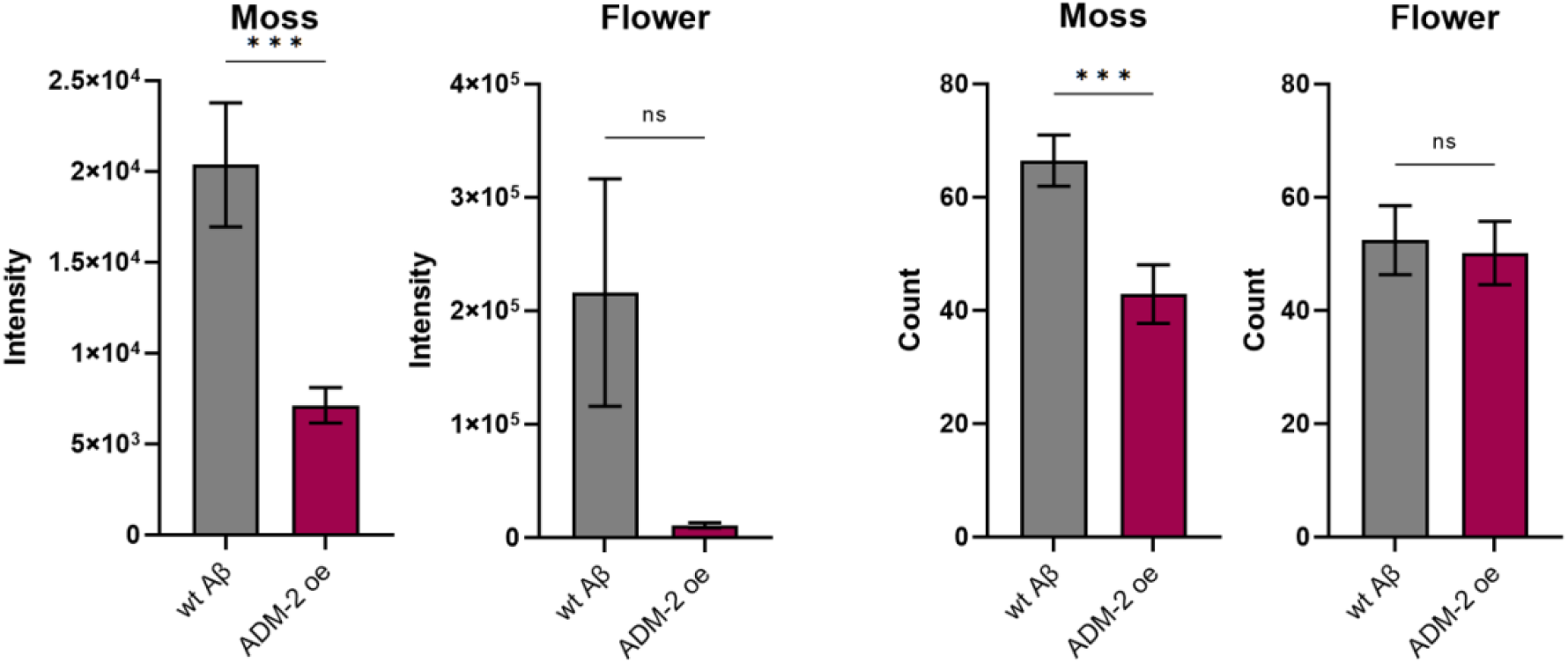
Overexpression of ADM-2 is sufficient to lead to a significant reduction of sfGFP::Aβ aggregates. Both the count and intensity of moss aggregates were reduced, but not of flower aggregates, when measured 24h after induction of ADM-2 overexpression. Data is the intensity and count measures over two independent experiments. Plotted are mean and SEM. Statistical analysis: unpaired *t*-test.

